# LC3C–ATG4D regulatory axis supports coronavirus replication organelle formation independently of canonical autophagy

**DOI:** 10.64898/2026.08.10.743971

**Authors:** Štěpánka Vlachová, Lucia Iovine, Valentina Marano, Elena Polishchuk, Michelle Cillo, Lorena Donnici, Pedro Machado, Paolo Swuec, Carmine Settembre, Paolo Grumati, Raffaele De Francesco, Lina Herhaus, Mirko Cortese

## Abstract

Coronaviruses hijack host membranes to assemble ER-derived double-membrane vesicles (DMVs) that shield viral RNA replication from the cell intrinsic surveillance. Although DMVs morphologically resemble autophagosomes, whether and how autophagy factors actively support their biogenesis has remained elusive. Here, we identify a non-canonical requirement for the autophagy protein LC3C in β-coronavirus replication. Loss of LC3s impaired viral RNA replication, whereas genetic ablation of ATG7 did not, indicating that canonical ATG7-dependent lipidation is dispensable in this context. Reconstitution experiments showed that only LC3C substantially restored replication in LC3-deficient cells and that LC3C phospho-mutants, differing in accessibility to ATG4-mediated processing, displayed distinct proviral activities. Additionally, ATG4D, the main protease responsible for maintaining the LC3 non-lipidated pool, is selectively required for viral replication. Both ATG4D and LC3s depletion triggers formation of aberrant DMV-like structures and potently suppresses SARS-CoV-2 replication. Ultrastructural analysis of nsp3–nsp4-induced membranes showed that depletion of LC3s or ATG4 proteases altered DMV abundance and morphology, supporting a role for the LC3C–ATG4D axis in replication organelle biogenesis. These data establish that β-coronaviruses repurpose ATG4D-driven LC3C de-lipidation for non-canonical LC3 recruitment to replication organelles, identifying the lipidation state of LC3 as a molecular determinant of replication organelle biogenesis and efficient viral replication.

**Highlights:** The manuscript shows that β-coronavirus replication depends on LC3 proteins and particularly on LC3C in reconstitution experiments, that this dependency is independent of ATG7-mediated lipidation, and that ATG4D promotes efficient replication and replication organelle morphology. Together, the data support a model in which a non-canonical LC3C–ATG4D pathway contributes to DMV biogenesis and viral RNA replication.

## Introduction

Positive-sense single-stranded RNA viruses replicate their genomes in association with virus-induced membranous replication organelles that spatially organise viral and host enzymes, concentrate metabolites, and shield replication intermediates from innate immune sensing. Coronaviruses induce extensive rearrangements of the endoplasmic reticulum (ER) to generate autophagosome-resembling double-membrane vesicles (DMVs) that serve as hubs for viral RNA synthesis(1–3). While viral non-structural proteins nsp3 and nsp4 are sufficient to drive DMVs biogenesis, increasing evidence suggests that host factors required for membrane-shaping and lipid mobilization might be co-opted to support replication organelle formation and function(4–6).

Autophagy has long been proposed to contribute to coronavirus replication, largely due to the morphological similarity between DMVs and autophagosomes and the re-localisation of autophagy markers during infection(1,7–10). However, genetic studies have yielded conflicting results, with many core autophagy genes proving dispensable for viral replication(11). While several SARS-CoV-2 proteins initiate the autophagy pathway activation, the complete autophagic flux is subsequently prevented by the viral proteins (6–8,12–15). This has led to the emerging view that coronaviruses exploit selective components of the autophagy machinery without engaging canonical degradative autophagy. Evidence suggests that SARS-CoV-2 exploits early autophagy components to support its replication, while blocking late steps to evade lysosomal clearance(6,7,12–14,16).

Central to autophagy is the ATG8 family of ubiquitin-like proteins, which in mammals comprises the LC3s (LC3A, LC3B, LC3C) and GABARAPs (GABARAP, GABARAP-L1, and GABARAP-L2) subfamilies. These proteins are usually processed by two ATG7-dependent conjugation systems to form an amide bond between the C-terminal carboxyl group of LC3s/GABARAPs and the amino group in the hydrophilic head of the phosphatidylethanolamine, a modification essential for LC3 proteins conjugation to autophagosomal membranes through the progression from inactive, cytoplasmic pro-LC3 to primed cytoplasmic LC3-I and finally membrane-bound lipidated LC3-II form(17,18). Lipidation is reversible and tightly regulated by four ATG4 cysteine proteases (ATG4A–D): ATG4B and ATG4A primarily prime newly synthesised ATG8 proteins by exposing their C-terminal glycine, whereas ATG4D and ATG4C preferentially delipidate membrane-associated forms(19– 22). Notably, LC3 paralogues exhibit distinct biochemical properties, expression patterns, and regulatory post-translational modifications, suggesting non-redundant cellular functions within and possibly beyond canonical autophagy(23–25).

Here, we investigate the contribution of LC3 proteins and their regulatory machinery to coronavirus replication. We identify LC3 proteins, and particularly LC3C in reconstitution experiments, as important host determinants of efficient replication of SARS-CoV-2 and HCoV-OC43. We further show that ATG7-mediated lipidation is dispensable, whereas the delipidating protease ATG4D promotes efficient replication and contributes to replication organelle morphology. These findings support a model in which β-coronaviruses repurpose a non-canonical LC3C–ATG4D pathway, distinct from canonical degradative autophagy, to support replication organelle biogenesis.

## Results

### An siRNA screen identifies ATG4D and LC3C as host dependency factors for coronavirus infection

To systematically dissect the contribution of autophagy-related genes to distinct stages of coronavirus infection, we performed a targeted siRNA screen against 48 core and regulatory ATGs. The screening followed a two-step approach to distinguish between factors affecting virus entry and replication from factors affecting assembly and release. In the primary screen (step 1), human pulmonary carcinoma A549 cells stably expressing ACE2 and TMPRSS2 were transfected with siRNAs for 48⍰h and subsequently infected with SARS-CoV-2 for 24⍰h. Next, supernatants from the primary infection were collected and used to infect naïve A549-ACE2-TMPRSS2 cells (step 2) (**Figure 1a**). Infection levels were quantified by high-content microscopy based on immunostaining for the viral nucleoprotein (N) and double-stranded RNA (dsRNA), a hallmark of active RNA virus replication. In parallel, LC3B staining was used as a control to monitor autophagy activation. As expected, N exhibited a diffuse cytoplasmic distribution, and cells positive for N were consistently positive for perinuclear dsRNA, which marks the viral replication organelles, whereas LC3B displayed both diffuse and punctate staining, reflecting cytosolic LC3-I and membrane-associated LC3-II, respectively (**Figure 1b**). Cell viability was assessed at the time of infection to exclude cytotoxic effects of gene silencing; only conditions maintaining >80% viability were included in downstream analyses (**Figure 1c, d**). Hits were defined based on significant changes in the percentage of infected cells (|z|⍰≥⍰1.5, p⍰< ⍰0.05) under conditions of acceptable viability (**Figure 1e, f**).

**Figure 1:**
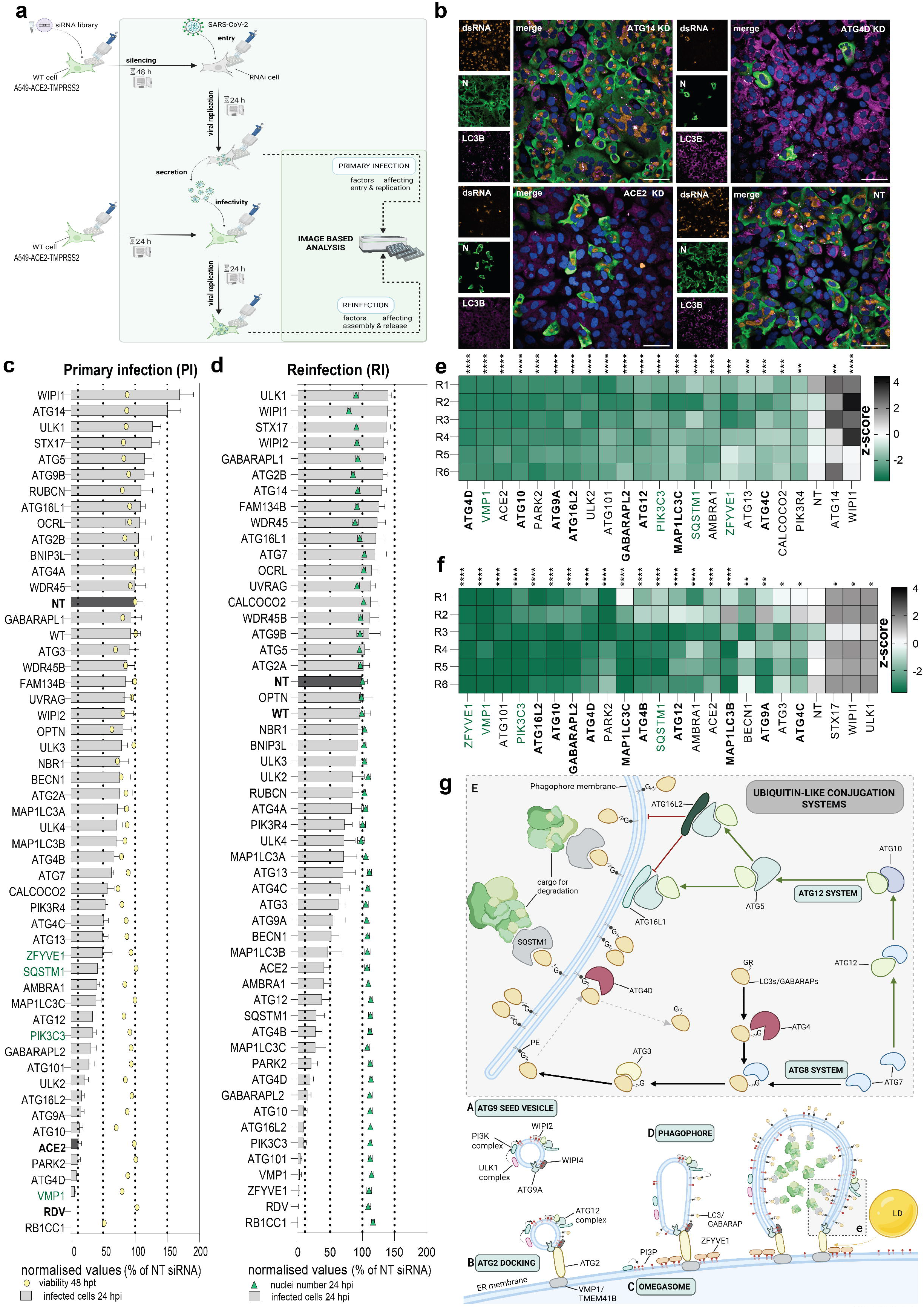
siRNA screening reveals novel autophagy factors affecting SARS-CoV-2 infection. **(a)** Schematic representation of screening experimental design – with the two-steps approach comprising of a primary infection of silenced cells and or reinfection of naive cells using virus containing supernatants derived from the primary screening. **(b)** Representative fluorescence microscopy images from high-content screening readout. Nuclei – Hoechst (blue), dsRNA (orange), SARS-CoV-2 Nucleocapsid (green), and LC3B (magenta). Insets show enlarged views of the boxed areas. Scale bar: 50 µm. **(c, d)** Percentage (%) of infection (grey bars) in primary infection **(c)** and reinfection **(d)** was quantified. Results are representative from two independent experiments with technical triplicates. Viability, quantified by ATP content at time of infection, is represented by yellow circle. Nuclei number at 24 hpi is represented by green triangle. Bar graphs represent the mean⍰fold change (±⍰SEM) of SARS-CoV-2 positive cells relative to NT cells. Remdesivir (RDV) was added as positive controls. Factors highlighted in green are already described dependency factors of SARS-CoV-2. **(e, f)** The z-score of individual factors in primary **(c)** and reinfection **(d)** screening were quantified and statistical significance was determined in GraphPad Prism version 10.5.0 using two-way ANOVA with Dunnett test (ns [not significant] p⍰>⍰0.05, *p⍰<⍰0.05, **p⍰<⍰0.01, ***p⍰<⍰0.001, **** p⍰<⍰0.0001). Factors were considered hits if they met the following criteria: p-value < 0.05, z-score ∣z∣≥1.5, and cell viability > 80%. Factors involved in Ub-like systems and phagophore expansion are highlighted in bold, previously reported host factors are shown in green. **(g)** Scheme of phagophore formation and involvement of Ub-like systems in autophagy.

This screening identified 19 dependency factors and 2 restriction factors, the majority of which have not previously been reported as SARS-CoV-2 host factors (**Figure 1e**). Notably, 43.8% of the targeted autophagy genes significantly affected infection, with 39.6% of perturbations leading to reduced viral replication, underscoring a prominent role for the autophagy machinery in supporting SARS-CoV-2 infection. The secondary reinfection screen largely confirmed the primary hits and further revealed factors with a possible stronger impact on virion assembly or release, including MAP1LC3B, ATG4B, ULK1, and STX17 (**Figure 1f**).

Strikingly, by clustering the screened factors into functional groups, we noted that majority of the dependency factors belonged to the ubiquitin-like conjugation system involved in early phagophore formation (**Figure 1g**). Among these, ATG16L2, which is a dominant-negative paralog of ATG16L1 capable of forming ATG5–ATG12 complexes that do not support canonical LC3 lipidation, emerged as a significant hit. ATG4D, a major delipidating protease for selected ATG8 family members including LC3C and GABARAPL2, emerged as the strongest dependency factor in the screen too.

### LC3 homologues are indispensable for β-coronavirus replication

The identification of LC3C among the strongest hits prompted us to investigate the requirements of LC3 homologues for SARS-CoV-2 infection. Consistently with previous reports, canonical autophagy marker LC3B accumulates in the membrane-bound lipidated form in our model system A549-ACE2-TMPRSS2 upon SARS-CoV-2 infection, while the subsequent autophagic flux is impaired as shown by SQSTM1 immunoblot levels (**Supp. Figure S1a**). Moreover, quantification of endogenous LC3B puncta in infected cells at 24hpi showed an increase in LC3B puncta number per cell compared to uninfected cells, where LC3B signal was mainly cytoplasmic (dashed line in **Supp. Figure S1b, c**), confirming that SARS-CoV-2 infection induces accumulation of LC3B.

To assess the requirement for LC3 proteins, triple LC3A/B/C knockout (KO) HeLa cells transiently transduced to express ACE2 and TMPRSS2 were infected with SARS-CoV-2. Quantification of infection by immunofluorescence at 48⍰h post-infection revealed a pronounced reduction (72%) in the percentage of virus-positive cells in LC3-deficient cells compared with wild-type controls (**Figure 2a**). This defect was accompanied by a ~90% decrease in intracellular viral RNA levels, indicating a severe impairment of viral replication (**Figure 2b**). No decrease on cell growth has been observed, confirming that the reduction in replication was not due to a metabolic defect (**Supp. Figure S1d**). Furthermore, to exclude general defects in gene expression, KO cells were transfected with a GFP-reporter expressing construct that showed comparable GFP intensity to WT (**Supp. Figure S1e**).

**Figure 2:**
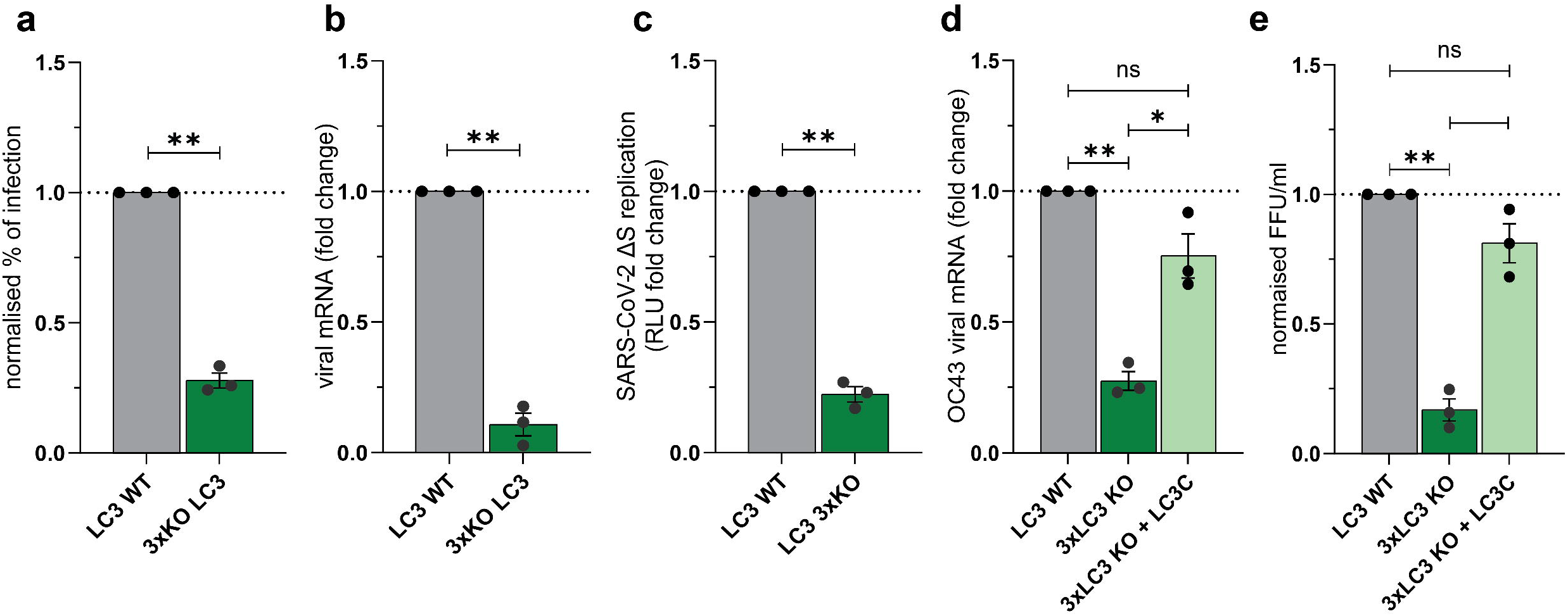
Lack of LC3 homologues impairs β-coronavirus infection. (**a**) HeLa knock-out for LC3A, B, and C (3xKO LC3) and their control cells were infected with moi=5 of SARS-CoV-2. Cells were fixed and permeabilised 48 hpi. Indirect immunofluorescence staining was performed to stain N protein and nuclei visualised by Hoechst. The percentage of infection was evaluated by number of N-positive cells. (**b**) 3xKO LC3 and control cells were infected with moi=5 of SARS-CoV-2. Viral subgenomic mRNA was quantified 24 hpi by q-RT-PCR. (**c**) 3xKO LC3 and control cells were transfected with 200ng of recombinant SARS-CoV-2-ΔS RLuc replicon. Cells were lysed and luminescence-based replication readout was performed 24 hpi. (**d-e**) 3xKO LC3, control and 3xLC3 KO reconstituted with LC3C were infected with moi=5 of HCoV-OC43 for 48 hours. Intracellular RNA was quantified by qRT-PCR using HCoV-OC43 ORF1 specific primers and TCID_50_ was performed to quantify secreted infectivity. Data shown in graphs are mean ± SEM from 3 independent experiments, each performed in technical replicates. Statistical significance was assessed using a one-sample t-test (ns [not significant] p⍰>⍰0.05, *p⍰<⍰0.05, **p⍰<⍰0.01, ***p⍰<⍰0.001, **** p⍰<⍰0.0001).

To directly determine the exact step of viral replication requiring LC3 homologues, we utilized a BAC-vectored SARS-CoV-2 replicon reporter, generated by deletion of the Spike glycoprotein (SARS-CoV-2 ΔS). Replicon activity was reduced by approximately 80% in LC3 triple KO cells relative to wild-type controls (**Figure 2c**), confirming that LC3 proteins act at a replication step independent of viral entry or particle assembly.

Interestingly, the requirement for LC3 homologues extends beyond SARS-CoV-2 since infection of LC3 triple KO cells with the human coronavirus OC43 resulted in a marked reduction in viral replication and progeny infectivity at 48⍰h post-infection (**Figure 2d, e**). Reintroduction of LC3C, the strongest hit among the ATG8 paralogues from the screening, in LC3 triple KO HeLa cells significantly restored OC43 replication and infectivity (**Figure 2e**). These findings identify LC3 proteins as conserved host factors required for efficient β-coronavirus replication and show that LC3C alone is sufficient to substantially restore replication in LC3-deficient cells, consistent with an important role for this paralogue during infection.

### SARS-CoV-2 replication proceeds independently of ATG7-mediated LC3s lipidation

Among the autophagy-related genes emerging from our screen were several components of the ATG8 conjugation machinery that participate in the two ubiquitin-like conjugation systems driving LC3/GABARAP lipidation. Notably, both conjugation cascades depend on the E1-like activating enzyme ATG7, which initiates the ATG8 and ATG12 pathways. Although ATG7 itself did not score as a hit in the screen, its central position in this machinery prompted us to investigate whether canonical LC3 lipidation is required for SARS-CoV-2 replication. To this end, we depleted ATG7 in the human lung adenocarcinoma Calu-3 cells and A549-ACE2 cells. While KD reduced ATG7 mRNA and protein levels, it did not affect viral replication in either cell line (**Figure 3a-f**).

**Figure 3:**
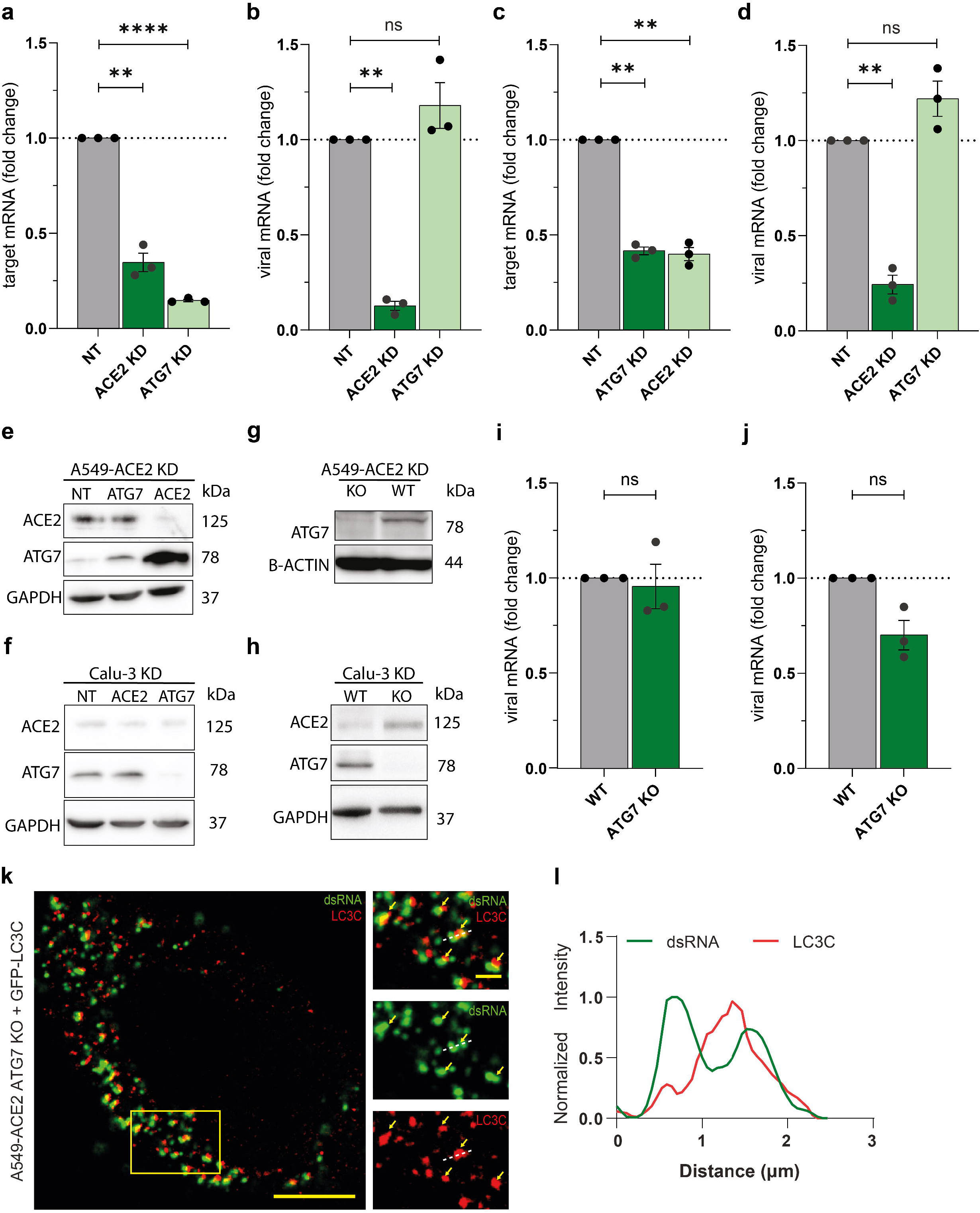
ATG7-mediated conjugation of ATG8 homologues is dispensable for the SARS-CoV-2 infection. (**a-b**) A549-ACE2 cells were transfected with non-targeting (NT), ATG7 or ACE2 siRNA (20 nM). Knockdown efficiency was assessed 48 hrs post-trasfection, after which cells were infected SARS-CoV-2 (MOI=1). Viral RNA levels were quantified 24 hpi by q-RT-PCR. (**c-d**) Calu-3 cells were transfected and infected as above and KD efficiency (**c**) and viral RNA (**d**) were quantified by q-RT-PCR. (**e-f**) Immunoblot of KD A549-ACE2 and Calu-3 cells. GAPDH served as loading control. (**g-h**) Immunoblot of ATG7 KO A549-ACE2 (**g**) and Calu-3 cells (**h**). β-actin and GAPDH served as loading control. (**i-j**) Viral RNA levels in ATG7 KO A549-ACE2 (**i**) and Calu-3 (**j**) cells infected with SARS-CoV-2 (MOI=1, 24 hpi). (**k**) Representative confocal images of A549-ACE2-ATG7 KO transfected with LC3C-GFP for 16 hours and infected with SARS-CoV-2 (MOI=5, 6 hpi). The cells were stained for dsRNA (green); LC3C-GFP is shown in red. Insets shows magnified boxed area; yellow arrows indicate dsRNA/LC3C double-positive puncta. Scale bars: 10⍰μm and 2 μm (main and inset, respectively). (**l**) Fluorescence intensity profile along the dashed line in (**k**). Intensities are shown normalized to the respective channel maximum. Data shown in graphs are mean ± s.e.m. from 3 independent experiments. Statistical significance was assessed using a one-sample t-test (ns [not significant] p⍰>⍰0.05, *p⍰<⍰0.05, **p⍰<⍰0.01, ***p⍰<⍰0.001, **** p⍰<⍰0.0001).

To exclude the role of ATG7-mediated lipidation process in SARS-CoV-2 infection, we generated CRISPR/Cas9 mediated knock-out (KO) of ATG7 in both cell lines. The ATG7 depleted cells were characterized by immunoblot of ATG7 and analysis of LC3B puncta upon nutrient starvation in A549-ACE2 ATG7 KO and WT cells (**Figure 3g, h, Supp. Figure S2a, b**). Consistently with expectations, LC3B positive foci were observed upon starvation of wild-type (atg7+/+) cells, but not in ATG7-deficient (atg7−/−) cells (**Supp. Figure S2a, b**). The ATG7 KO cells were then infected with SARS-CoV-2 and subgenomic viral mRNA levels and viral titers were analysed at 24 hpi. In agreement with previous results, viral replication was not affected by KO of ATG7 protein (**Figure 3i, j**), demonstrating that canonical ATG7-dependent LC3 lipidation is dispensable for SARS-CoV-2 replication.

Although no autophagic puncta could form in ATG7KO cells upon starvation, in line with lipidation impairment, numerous GFP-tagged LC3C foci were observed at early time post-infection in close proximity of dsRNA, suggesting that the replication impairment observed upon LC3-KO can be due to the need of non-lipidated, possibly recycled, LC3C homologues, such as those regulated by ATG4D (**Figure 3k-l**). Together, these results indicate that SARS-CoV-2 replication does not require ATG7-dependent LC3 lipidation and support a non-canonical role for LC3 proteins in coronavirus replication distinct from canonical ATG7-dependent autophagy.

### ATG4D is required for efficient replication of β-coronaviruses

After confirming that lipidation of LC3s is not required for the infection, we next sought to examine the role of ATG4D, which is a prominent host dependency factor from the screening and the main LC3s delipidating enzyme. We investigated its role in SARS-CoV-2 infection at a mechanistic level. Notably, previous work demonstrated that ATG4D preferentially regulates the recycling of LC3C and GABARAPL2, establishing a direct functional axis between ATG4 delipidating activity and these ATG8 family members(22,26).

First, to validate the screening results, ATG4D was depleted using an independent set of siRNAs, and the analysis was extended to human kidney proximal tubule (HK2) cells to assess whether the observed phenotype was cell type dependent. HK2 cells were transfected with siRNAs for 48⍰h, after which cell viability was assessed prior to infection with SARS-CoV-2 (MO⍰= ⍰1) for 24⍰h. Knockdown efficiency and viral replication were quantified by qRT–PCR of ATG4D mRNA and viral sub genomic RNA levels, respectively. Reduction of ATG4D mRNA levels was associated with a decrease in viral RNA and secreted infectivity in both A549-ACE2 and HK2 cells (**Figure 4a-c** and **Supp. Figure S3a-c**).

**Figure 4:**
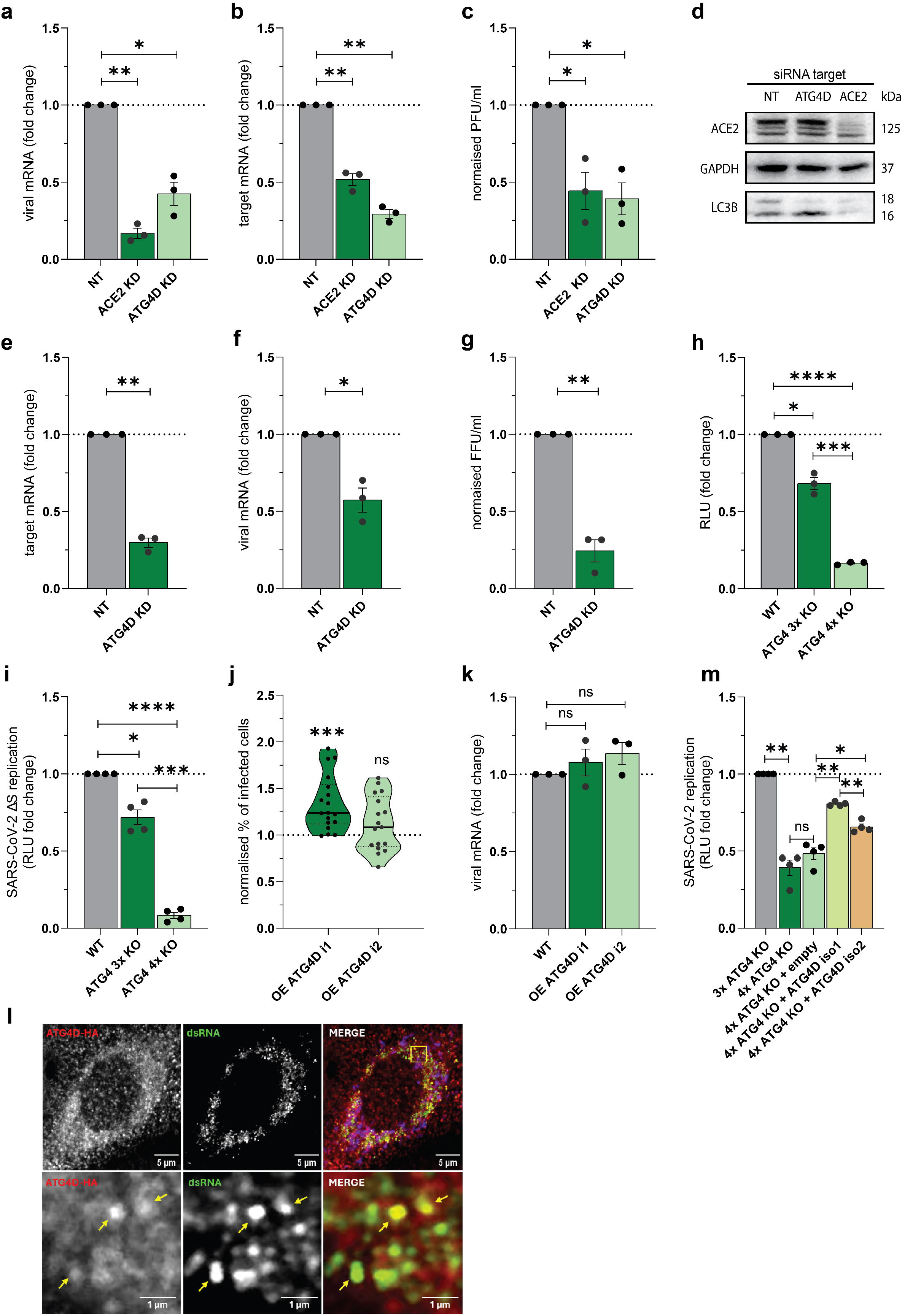
ATG4D knock-down reduces SARS-CoV-2 infection. (**a–c**) A549-ACE2 cells were transfected for 48 h with ATG4D, ACE2 or NT siRNA (20 nM) and infected with SARS-CoV-2 (MOI=1). After 24 hpi, viral replication (**a**) and knockdown efficiency (**b**) were quantified by q-RT-PCR and extracellular infectivity was quantified by plaque assay (**c**). (**d**) Immunoblot of ATG4D-silenced A549-ACE2 cells at 48 hpt. GAPDH served as loading control. **(e–g)** MRC5 cells were transfected for 48 h with NT or ATG4D siRNA (20 nM) and infected with HCoV-OC43 (MOI=0.05). After 24 h, ATG4D knockdown efficiency (**e**) and intracellular viral ORF1a mRNA levels (**f**) were quantified by RT-qPCR, while extracellular infectivity was quantified by TCID_50_ (**g**) on Vero E6 cells. (**h**) HeLa WT, ATG4 3x KO (A/B/C) and ATG4 4x KO (A/B/C/D) expressing ACE2-TMPRSS2 were infected with recombinant SARS-CoV-2-RLuc (MOI=1) and replication was assessed by luminescence quantification 24 hpi. (**i**) HeLa WT, ATG4 3x KO and ATG4 4x KO were transfected with SARS-CoV-2-ΔS RLuc BAC construct. Luminescence was quantified 24 hpi. **(j–k)** A549-ACE2-TMPRSS2 cells stably overexpressing HA-ATG4D isoform 1 or 2 were infected with SARS-CoV-2 (MOI=1). After 24 hpi, the percentage of N-positive cells **(j)** and subgenomic mRNA levels **(k)** were quantified. (**l**) Representative confocal images of SARS-CoV-2-infected A549-ACE2-TMPRSS2 cells overexpressing HA-ATG4D isoform 1, stained for HA (red) and dsRNA (green). Yellow arrows indicate colocalising signal. (**m**) HeLa ATG4 triple and tetra KO cells transduced with ACE2-TMPRSS2 and reconstituted with empty vector, ATG4D isoform 1 or isoform 2 were infected with SARS-CoV-2-RLuc (MOI=1) and replication assessed by luminescence at 24 hpi. Data are mean ± s.e.m. from ≥3 independent experiments in technical triplicates. Statistical significance assessed by one-sample t-test against NT or WT (theoretical mean = 1; ns p > 0.05, *p < 0.05, **p < 0.01, ***p < 0.001, ****p < 0.0001).

Despite multiple attempts, endogenous ATG4D could not be reliably detected using available commercial antibodies. However, functional validation of KD was performed by immunoblot analysis that showed increased levels of LC3B-II upon ATG4D knockdown in both A549-ACE2 and HK2 cells, in line with ATG4D being the main LC3-II delipidating enzyme (**Figure 4d, and Supp. Figure S3d**).

To assess whether the requirement for ATG4D extends to other β-coronaviruses, human lung fibroblast cells MRC5 were depleted of ATG4D by siRNA-mediated knockdown and subsequently infected with HCoV-OC43. At 24 hpi, both replication and infectivity were significantly reduced (**Figure 4e-g**). These findings indicate that LC3 delipidation mediated by ATG4D appears to be a conserved prerequisite for efficient β-coronavirus replication.

In humans, the ATG4 protease family comprises four homologues—ATG4A, ATG4B, ATG4C, and ATG4D—which share a conserved catalytic cysteine protease domain but differ substantially in their N- and C-terminal regions. Although partial functional redundancy has been reported, these proteases display distinct substrate specificities and differential priming and deconjugation activities toward ATG8 family members. ATG4B is responsible for the majority of ATG8 priming events and can also mediate ATG8 delipidation, albeit much less efficiently than ATG4D. In contrast, ATG4D exhibits relatively weak priming activity but functions as a major delipidating enzyme for specific ATG8 paralogs, including LC3C and GABARAPL2.

To exclude compensatory effects among ATG4 homologues, we compared HeLa wild-type (WT) cells with isogenic lines lacking either three ATG4 homologues (3×ATG4 KO, retaining only ATG4D) or all four of them (4×ATG4 KO). To first rule out any general cellular defects introduced by the knockouts, we assessed episomal GFP expression across cell lines. Compared to WT, 4×ATG4 KO cells showed a 17% reduction in GFP intensity and 3×ATG4 KO cells a 29% reduction, indicating a moderate impact on protein expression in both knockout backgrounds (**Supp. Figure S3e**). Next, we infected all cell lines with a recombinant SARS-CoV-2 expressing Renilla luciferase and quantified the Renilla activity as proxy for viral replication at 24 hours post-infection (MOI = 1). Complete loss of ATG4 activity in 4×ATG4 KO cells resulted in a marked reduction of Renilla compared to WT (**Figure 4h**). Retention of endogenous ATG4D alone, as in 3×ATG4 KO cells, was sufficient to rescue the defect in infection and viral replication, demonstrating a profound role of sole ATG4D protein in supporting SARS-CoV-2 replication (**Figure 4h**).

To directly assess whether ATG4 proteases are required at the level of viral RNA replication, we employed a bacterial artificial chromosome (BAC)–vectored SARS-CoV-2 ΔS replicon encoding Renilla luciferase as a reporter. HeLa wild-type, 3×ATG4 KO, and 4×ATG4 KO cells were transfected with the replicon for 24⍰h, and luciferase activity was measured as a readout of replication. Consistent with infection-based assays, loss of ATG4A/B/C resulted in a modest reduction in replicon activity, whereas deletion of all ATG4 homologues caused a strong 92% decrease in replication (**Figure 4i**). These data confirm that ATG4 protease activity, potentially dominantly mediated by ATG4D, is specifically required for efficient SARS-CoV-2 RNA replication.

ATG4D exists in two isoforms with distinct subcellular localizations, with a longer cytoplasmic isoform 1 and a shorter isoform 2 localized to mitochondria (**Supp. Figure S3f**). To dissect their individual contributions to viral infection, we transduced A549 cells with lentiviruses encoding an HA-tagged codon-optimized ATG4D to achieve stable protein overexpression (OE). To assure comparable results between ATG4D isoforms we transduced various moi of ATG4 lentiviruses and selected the ones expressing similar protein amount (**Supp. Figure S3g**). As reported in literature, immunofluorescence staining of the longer ATG4D isoform 1 displayed a homogenous cytoplasmic distribution and only in a minority of cells shows mitochondrial localisation, which occurs only after caspase 3 cleavage in DEVD-K motif (**Supp. Figure S3h**). Consistently with its predicted mitochondrial localization, the shorter isoform 2, created by alternative splicing, extensively colocalized with mitochondria, as shown by the colocalization with the mitochondrial marker TOMM20 (**Supp. Figure S3i**).

Both the ATG4D OE cells and the empty vector control cells were transduced with ACE2-TMPRSS2 lentivirus and subsequently infected with moi=1 of SARS-CoV-2 for 24 hours. Cells overexpressing isoform 1 exhibited a modest increase in the percentage of infected cells compared to controls (**Figure 4j**), however, no significant changes were observed in viral sub genomic mRNA, suggesting that endogenous levels of ATG4D are sufficient for optimal SARS-CoV-2 replication and ATG4D is therefore not the limiting factor for infection under these conditions (**Figure 4k**). Interestingly, upon immunofluorescent staining of ATG4D in the infection context we observed recruitment of HA-tagged ATG4D isoform 1 to the area of dsRNA marking the vROs, while ATG4D isoform 2 show no noticeable changes (**Figure 4l, Supp. Figure S3j**). These data together with our previous observations suggest that the cytoplasmic isoform of ATG4D is the one preferentially hijacked by the virus and potentially contributing to viral infection.

To confirm the individual contributions of both isoforms to viral replication, we performed isoform-specific reconstitution experiments in 4×ATG4 KO cells, followed by infection with luciferase-expressing SARS-CoV-2 (MOI = 1). Expression of either isoform substantially restored viral replication to levels comparable to those observed in 3×ATG4 KO cells expressing endogenous ATG4D, with isoform 1 conferring a greater rescue than the mitochondria-targeted isoform 2 (80% and 69% of the control, respectively) (**Figure 4m**). These results indicate that while both ATG4D isoforms support SARS-CoV-2 replication the proviral activity of ATG4D is primarily exerted in the cytoplasm thanks to isoform 1.

### ATG4D-accessible LC3C supports coronavirus replication

To test whether viral replication is influenced by the ATG4-mediated proteolytic processing of LC3C, we reconstituted LC3C expression in triple LC3 knockout cells using phospho-mimetic mutants designed to mimic either the phosphorylated (S93/96D) ATG4-inaccessible proLC3C or the non-phosphorylated (S93/96A) ATG4-accessible form of the LC3C as previously described(26) (**Figure 5a**). This setup enabled us to determine whether LC3C’s role in viral infection depends on its conjugation status or on the general accessibility of its C-terminal tail to ATG4.

**Figure 5:**
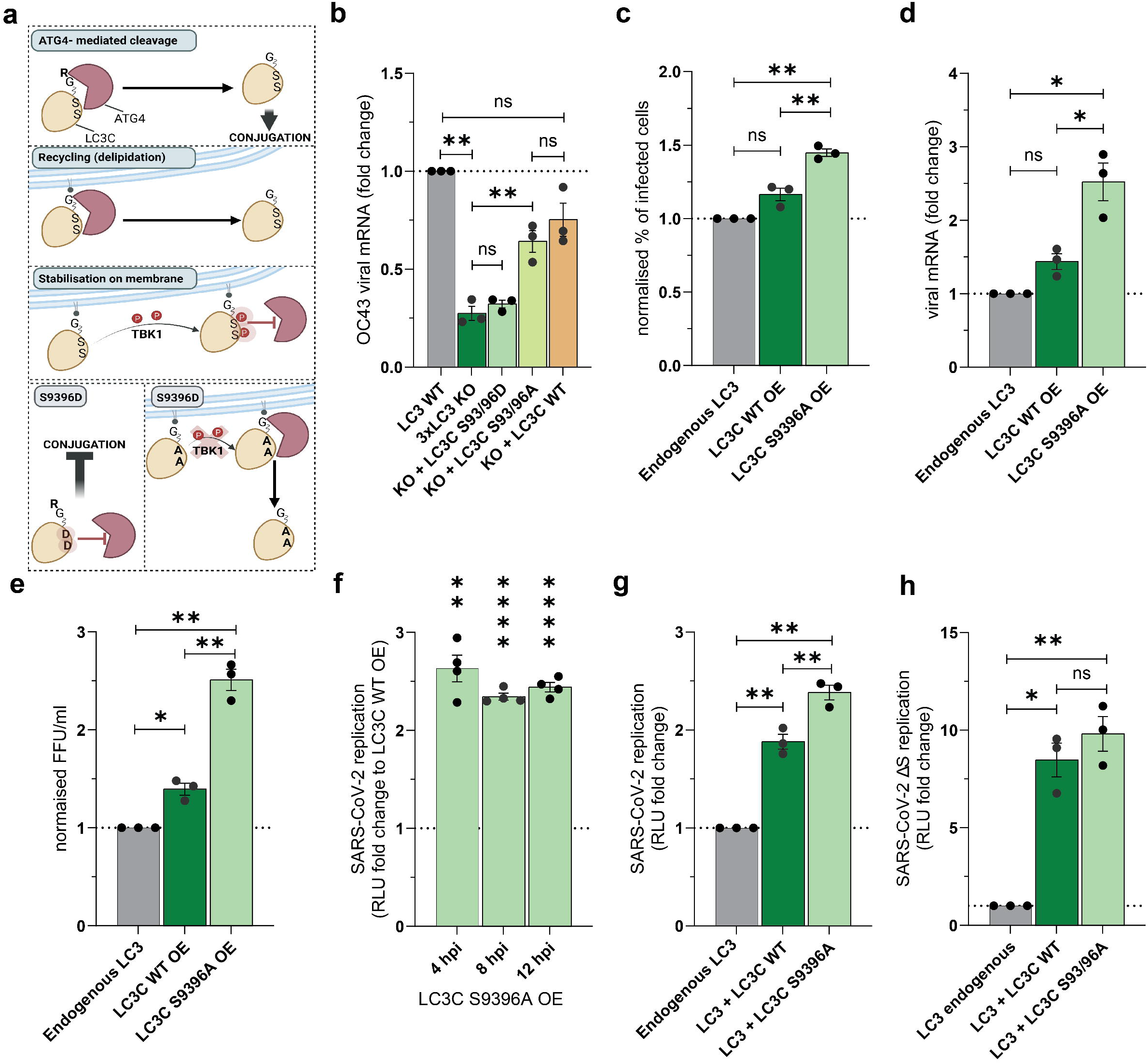
LC3C is limiting factor in β-coronavirus infection. (**a**) Schematic of ATG4-mediated LC3C processing. (**b**) HeLa LC3 3xKO cells were transfected with the indicated constructs and, after 4 hours, infected with HCoV-OC43 (MOI=5). Intracellular viral RNA was extracted and quantified by q-RT-PCR 48 hpt. (**c-e**) U2OS WT or overexpressing eGFP-mCherry-LC3C WT and S93/96A mutant were infected with HCoV-OC43 (MOI=5). After 24 hrs, percentage of infected cells (c), intracellular viral RNA (**d**) and extracellular infectivity (**e**) were quantified. (**f**) U2OS cells overexpressing LC3C S93/96A were infected with Rluc-SARS-CoV-2 (MOI=5) and luminescence was quantified 4, 8 and 12 hpi. (**g**) U2OS WT, U2OS-eGFP-mCherry-LC3C WT and LC3C S93/96A overexpressing cells were infected with Rluc-SARS-CoV-2 (MOI=5). Viral replication was assessed by luminescence at 8 hpi. (**h**) U2OS WT, U2OS-eGFP-mCherry-LC3C WT and LC3C S93/96A cells were transfected with recombinant SARS-CoV-2-ΔS – RLuc BAC construct and luminescence was quantified 24 hpt. Data are mean ± s.e.m. from 3 independent experiments in technical replicates. One-sample t-test against theoretical mean of 1 (ns p > 0.05, *p < 0.05, **p < 0.01, ***p < 0.001, ****p < 0.0001).

HeLa 3xLC3 KO reconstituted with LC3C WT and phospho-mutants were infected with HCoV-OC43. Reconstitution with LC3C WT restored viral replication in LC3-deficient cells, as expected. Notably, the S93/96A mutant rescued replication to a comparable level of WT (**Figure 5b**). In contrast, the S93/96D variant did not enhance viral replication. The inability of the phospho-mimetic mutant to rescue infection indicates that ATG4D-mediated cleavage of LC3C into LC3C-I is essential for its proviral function.

To further validate the role of LC3C in supporting viral replication, we evaluated the effect of LC3C WT and phospho-mutant overexpression on coronavirus replication. To this aim, we utilized U2OS cell lines stably overexpressing eGFP–mCherry-tagged LC3C WT or the S93/96A mutant, sorted for comparable LC3C expression levels (**Supp. Figure S4a**). Upon HCoV-OC43 infection, overexpression of LC3C WT increased the percentage of infected cells, intracellular viral RNA, and production of infectious particles relative to parental cells. Strikingly, expression of the S93/96A mutant consistently enhanced replication beyond that observed with LC3C WT, suggesting that increasing the pool of ATG4-accessible LC3C potentiates viral infection (**Figure 5c-e**).

To determine whether this effect extends to SARS-CoV-2 infection, and to specifically assess its impact on early stages of the viral life cycle, we performed short-term infection kinetics with recombinant luciferase-expressing SARS-CoV-2-RLuc evaluating the replication at 4-, 8- and 12-hours post-infection, thereby minimizing potential confounding effects from viral assembly or spread. Remarkably, already at 4 hpi a 2.6-fold increase of viral replication was observed in the cells with LC3C S93/96A OE compared to the LC3C WT OE and this trend was consistent during the time course (**Figure 5f**). Interestingly, in SARS-CoV-2 infected cells, even the overexpression of LC3C WT alone enhanced replication to 1.88-fold (**Figure 5g**) compared to the endogenous LC3 levels, suggesting that increasing levels of LC3C have a proviral effect. These observations were further corroborated upon transfection of the SARS-CoV-2 BAC-ΔS system, where we observed 8.5-fold and 9.8-fold increase in replication in LC3C WT and LC3C S93/96A mutant, respectively. (**Figure 5h**).

The inability of the phospho-mimetic mutant to rescue infection indicates that access of the LC3C C-terminus to ATG4-mediated processing is important for its proviral activity. Together with the enhanced activity of the S93/96A mutant in overexpression settings, our data support a model in which LC3C species permissive for ATG4 processing promote coronavirus replication.

### LC3s and ATG4D are required for DMVs biogenesis

Given that both LC3C and ATG4D specifically promote viral replication and that their effect maps to an early stage of the replication cycle, we sought to determine whether they contribute to DMV formation. To this aim, we performed transmission electron microscopy (TEM) following ectopic expression of SARS-CoV-2 nsp3 and nsp4, which is sufficient to induce double-membrane vesicle (DMV) formation in LC3s and ATG4s deficient cell lines.

In control HeLa cells, nsp3–nsp4 expression resulted in abundant, spherical DMVs with closed double membranes, consistent with previously described replication organelle morphology. In contrast, triple LC3A/B/C knockout cells exhibited a significant reduction in the number of nsp3–nsp4–induced DMVs, and the structures that did form displayed pronounced morphological defects (**Figure 6a, b**). Rather than closed vesicles, DMVs in LC3-deficient cells were frequently open, with incomplete double membranes and significantly shorter membrane arcs, possibly indicative of impaired membrane elongation (**Figure 6c, d**). These observations suggest that LC3 family proteins collectively contribute to the efficient nucleation and membrane elongation of coronavirus-induced replication organelles.

**Figure 6:**
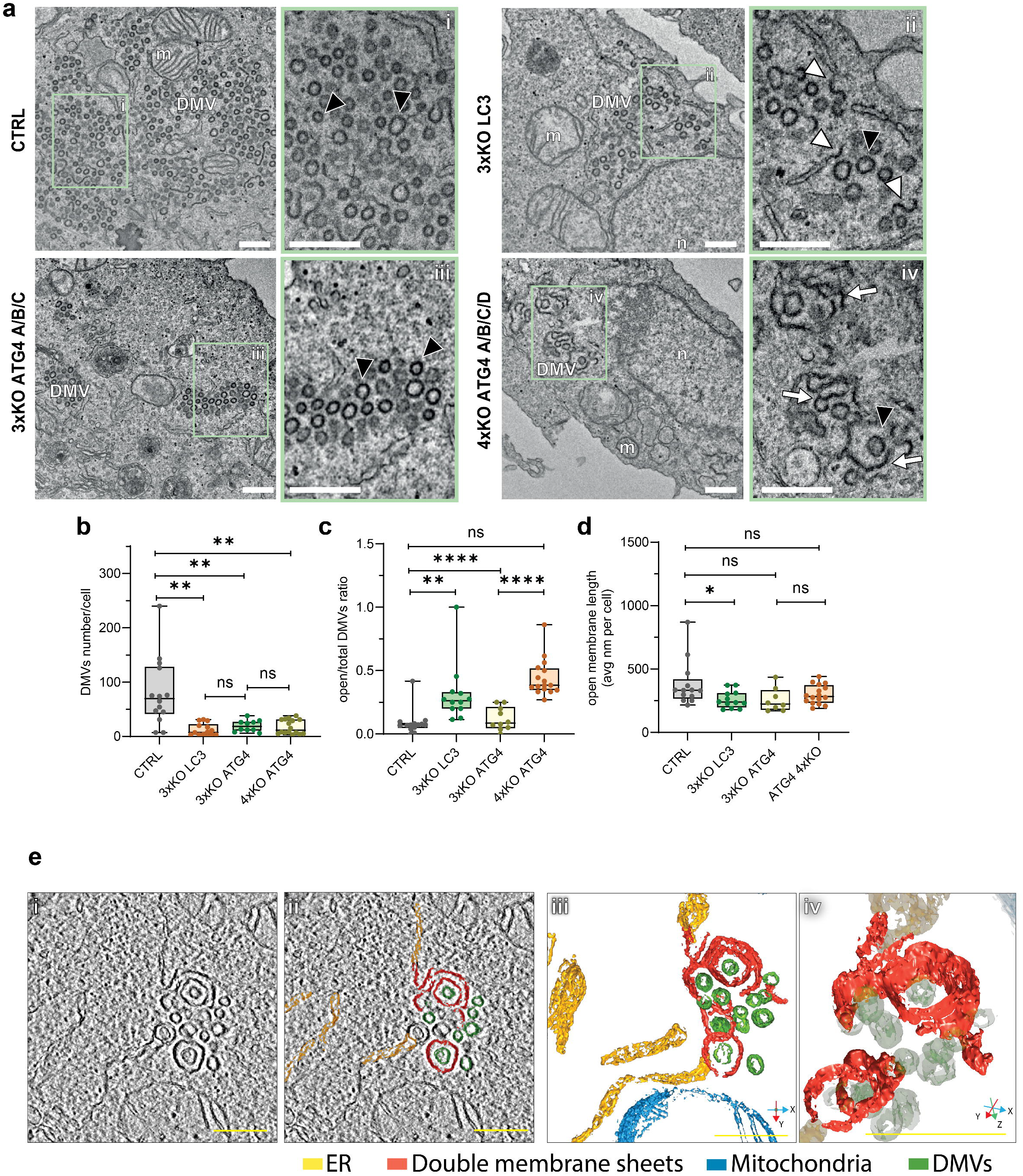
LC3 and ATG4D depletion alters DMV morphology. HeLa CTRL, 3xKO LC3, 3xKO ATG4 and 4xKO ATG4 cells were transfected with SARS-CoV-2 GFP-nsp3-nsp4-V5 polyprotein for 16 hours. (**a**) Representative TEM images of DMVs produced in HeLa CTRL, 3xKO LC3, 3xKO ATG4 and 4xKO ATG4 cells. Insets show morphological variations marked by arrows. Black arrowheads: closed DMV; white arrowheads: short open double membrane; white arrows: long open double membrane. Nuclei and mitochondria are labelled (n and m, respectively). (**b**) Quantification of DMV number/cell. (**c**) Quantification of open DMVs ratio compared to total DMVs. (**d**) Quantification of open double-membrane length/cell. (**e**) Electron tomography and 3D reconstruction of 4xKO ATG4 expressing GFP-nsp3-nsp4-V5 polyprotein. (i) Slice through the tomogram. (ii) Same region as in (i) with superimposed rendering of cellular and viral organelles that are specified on the bottom of the figure. (iii) 3D rendering of organelles visualized in (ii). Scale bars, 500 nm. See also Video S1.

Similarly, in cells lacking all four ATG4 paralogues displayed markedly impaired nsp3–nsp4-induced DMV biogenesis, with reduced DMV numbers and an accumulation of open, incompletely sealed double-membrane structures (**Figure 6b, c**). When observed in 3D, these structures resemble long sheets of double membrane that eventually bend and, in some cases seal, to form large double membrane vesicles, often surrounding properly formed DMVs (Figure 6e and Video S1). Retention of endogenous ATG4D in 3×ATG4 knockout cells restored DMV morphology toward that observed in control cells, although interpretation of DMV abundance in this background should consider the general reduction in protein expression noted during cell line characterisation (**Supp. Figure S3e**). Strikingly, selective removal of ATG4A, ATG4B, and ATG4C—while retaining endogenous level of ATG4D—restored DMV morphology, yielding closed vesicles comparable in appearance to those observed in control cells (**Figure 6a-c**). The selective retention of ATG4D restoring DMV closure, identifies ATG4D as the critical determinant of membrane sealing, implying its possible functional role in membrane scission and curvature stabilisation, likely through maintaining high pool of LC3-I through its recycling. Together, these data support a model in which LC3 proteins contribute to DMV membrane expansion, whereas ATG4D-dependent ATG8s processing promotes efficient maturation or closure of replication organelles (**Figure7**).

**Figure 7:**
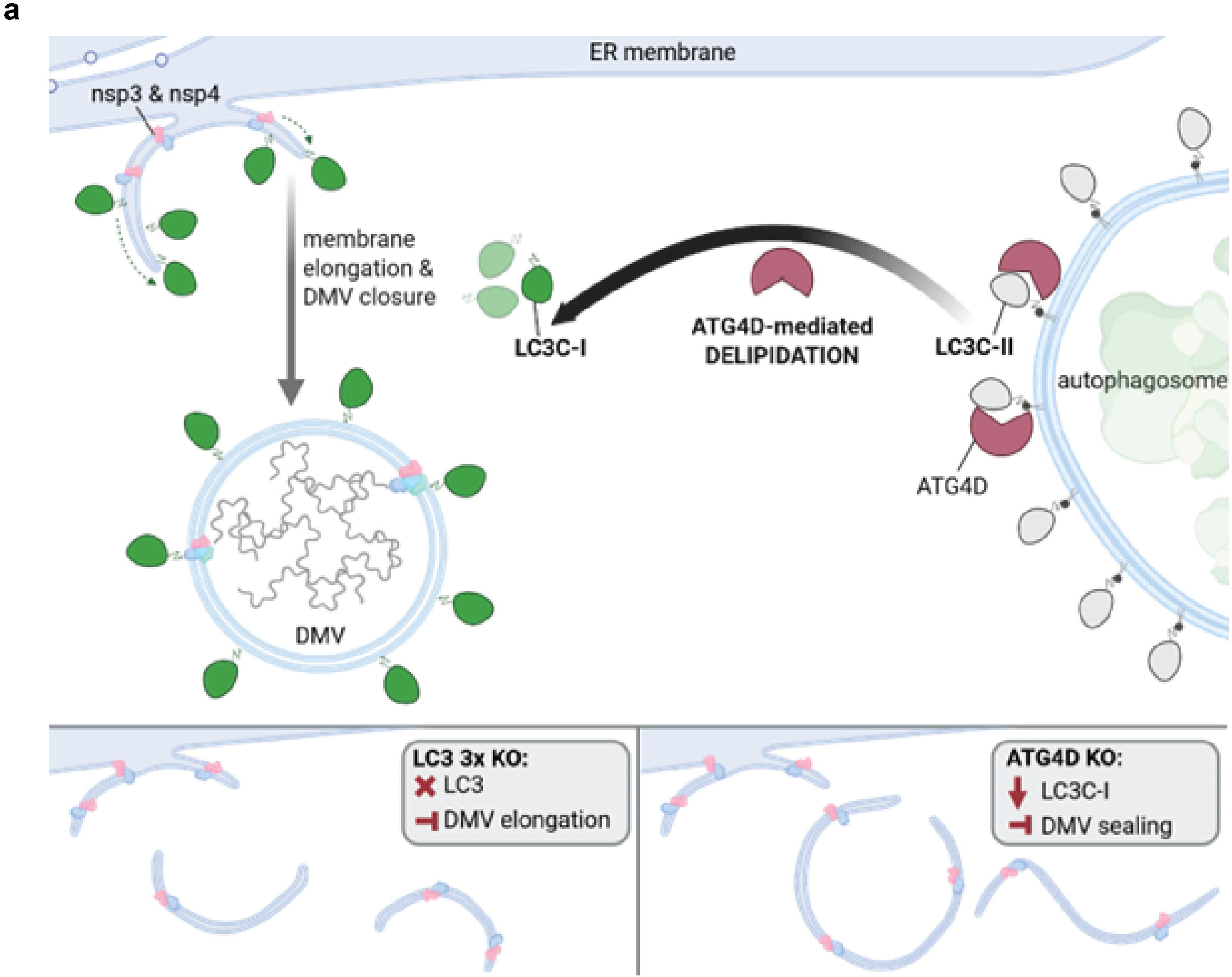
Model of LC3C and ATG4D function in SARS-CoV-2 DMV biogenesis. During coronavirus infection, nsp3 and nsp4 initiate ER membrane remodelling, driving membrane elongation and DMV closure. LC3C-I, recruited to the nascent DMV membrane, promotes membrane elongation and sealing. ATG4D-mediated delipidation of LC3C from the sealed DMV releases LC3C-I back to the cytosol, mirroring the LC3 cycle on autophagosomes. In the absence of LC3 (3xKO), membrane elongation is impaired, resulting in truncated open double-membrane structures. In the absence of ATG4D (KO), LC3C-I levels are reduced and DMV sealing is inhibited, leading to accumulation of unsealed double-membrane intermediates.

## Discussion

During coronaviruses replication, viral proteins and co-opted host-factors induce extensive remodelling of intracellular membranes to generate double-membrane vesicles (DMVs) that serve as replication organelles and shield viral RNA replication intermediates from host surveillance(4,6,7,27). Autophagy-related proteins, such as PIK3C3, ZFYVE1 or VMP1, have been implicated in coronaviral replication organelles biogenesis process(6,7,28). SARS-CoV-2 infection leads to the accumulation of LC3-positive membranes and disruption of autophagy flux to facilitate virion release, yet genetic perturbation of core autophagy components such as ATG5 doesn‘t influence viral replication(7–9,11–16,29–31).

Here, we establish that LC3 homologues are required for efficient β-coronavirus replication and uncover a non-canonical regulatory function of the ATG8s system in β-coronavirus replication that diverges from classical autophagy. Through systematic genetic interrogation, we showed that SARS-CoV-2 replication is profoundly impaired in LC3-deficient cells, whereas ATG7-dependent lipidation is dispensable. The lack of a replication phenotype upon ATG7 depletion, despite the pronounced defect observed in LC3-deficient cells, indicates that LC3 function in this context is independent of its conjugation to phosphatidylethanolamine. This requirement is not restricted to SARS-CoV-2, as HCoV-OC43 replication is similarly compromised in LC3-deficient cells, indicating that exploitation of LC3 proteins represents a conserved feature of β-coronavirus infection. These findings provide a framework to clarify why coronaviruses activate autophagy without the actual need of complete degradative flux *per se* and support a model in which LC3 proteins are repurposed outside canonical autophagy during infection.

Among the LC3 paralogues, LC3C emerged as a principal determinant of viral replication. Reconstitution experiments demonstrated that LC3C alone is sufficient to substantially restore replication in LC3-deficient cells, indicating that this paralogue can functionally compensate for the loss of other LC3 family members in the context of infection. LC3C has previously been associated with early phagophore assembly sites through selective interactions with initiation complex components ATG13, FIP200 and ATG9, events that precede and are spatially distinct from canonical LC3B recruitment, suggesting that unique structural properties of LC3C may be selectively exploited in the context of viral replication organelle formation(24).

Our data further indicate that the proviral function of LC3C depends on its accessibility to ATG4-mediated processing. TBK1-mediated phosphorylation of LC3C at residues S93 and S96 controls its accessibility to ATG4D, retaining LC3C in a lipidated membrane-associated form and preventing its delipidation and consequent return to the cytosolic non-lipidated pool. Phospho-mimetic mutations S93/96D in the LC3C C-terminal region that impair ATG4 cleavage abolished LC3C‘s ability to rescue replication, whereas phosphorylation-deficient S93/96A variant enhanced replication in comparison to wild-type LC3C. Given that these variants differ primarily in their susceptibility to ATG4-mediated proteolysis, these findings suggest that efficient LC3C turnover is required for its function during infection(26). Intriguingly, SARS-CoV-2 actively suppresses TBK1 activity through multiple viral proteins including M, NSP6 and NSP13, a strategy primarily described in the context of innate immune evasion. Considering our findings, this viral interference with TBK1 may carry an additional consequence: by reducing TBK1-mediated phosphorylation of LC3C at S93/S96, the virus would favour retention of LC3C in its non-lipidated, ATG4D-accessible form, thereby enhancing its own replication by potentiating the proviral ATG4D–LC3C axis(32).

While the four mammalian ATG4 proteases share conserved catalytic domains, our genetic analysis demonstrates that ATG4D, the strongest delipidating protease among ATG4s, displays the largest contribution to efficient coronavirus replication. Combined quadruple deletion of ATG4 paralogues resulted in a pronounced replication defect that was significantly rescued by selective retention of endogenous ATG4D, whereas loss of ATG4A, ATG4B, and ATG4C alone has a comparatively milder effect. In line with its established role as a delipidating enzyme, depletion of ATG4D leads to accumulation of lipidated LC3 and reduced viral replication, supporting a model in which ATG4D maintains a pool of cleaved LC3 available for function during β-coronavirus infection.

Ultrastructural analyses further provide mechanistic insight into the contribution of LC3C-ATG4D regulatory pathway to replication organelle biogenesis. Loss of LC3 proteins reduced the number of nsp3–nsp4-induced DMVs and was associated with the formation of shorter, incompletely sealed membrane structures, consistent with a defect in membrane elongation and stabilisation(33,34). In contrast, selective loss of ATG4D led to the accumulation of double membrane structures resembling an open DMVs, indicating impaired membrane closure. Further support for this model comes from electron tomography, which revealed that the double membrane structures observed correspond to large double membrane sheets that eventually bend and seal into aberrantly enlarged DMVs. Together, these observations support a model in which LC3 proteins contribute to membrane expansion, whereas ATG4D function regulates the maturation and sealing of replication organelles(33–35). In the absence of ATG4D, the double membrane continues to expand, forming a large sheet that eventually closes, giving rise to aberrantly large DMVs.

The preferential use of non-lipidated LC3C-I observed in our study is consistent with earlier work demonstrating that coronaviruses can recruit LC3 independently of canonical autophagy (36,37). Mouse hepatitis virus replication complexes associate with LC3B-positive membranes in an ATG7-independent manner, implicating LC3B-I rather than LC3B-II is present at coronavirus-induced replication structures(36). Our study extends these observations showing a broader and conserved mechanism among coronaviruses identifying LC3C as a relevant paralogue in human β-coronavirus infection and ATG4D as a critical regulator of LC3 processing, thus linking the availability of LC3C-I pool to replication organelle maturation and DMV closure.

Given the structural similarities of replication organelles across coronaviruses, it is plausible that LC3C-mediated membrane modulation provides a common platform for DMV biogenesis.

Taken together, our data support a model in which ATG4D-mediated proteolysis sustains a dynamic pool of cleaved, non-lipidated LC3C-I that supports the membrane remodelling and fusion events underlying DMV biogenesis. Rather than acting as a lipidated membrane anchor, LC3C-I may function as a scaffold that stabilizes highly curved double membranes or coordinates their remodelling. This lipidation-independent role is consistent with ATG7 being dispensable while interference with LC3C processing or ATG4D activity is detrimental, and would allow the virus to harness LC3-dependent membrane remodelling without engaging the degradative arm of autophagy. Defining how LC3C-I engages viral and host membrane-shaping factors will be key to understanding coronavirus membrane remodelling and the interplay between autophagy-related factors and viral replication. Overall, our findings support a conserved, non-canonical LC3C–ATG4D pathway that promotes replication organelle biogenesis and efficient viral RNA replication.

### Limitations of the study

A limitation of this study is that endogenous ATG4D could not be robustly monitored with available antibodies, which limited direct assessment of its localisation and processing dynamics during infection. In addition, structural conclusions from the nsp3–nsp4 system should be interpreted as mechanistic support for replication organelle biogenesis rather than a complete reconstruction of the infected-cell context. In this study we did not directly test whether the RO-forming viral proteins nsp3 and nsp4 physically engage LC3C or ATG4D, and neither factor was recovered among the nsp3–nsp4 interactors in previous proteomic characterization of the RO-forming polyprotein system (5). However, such interaction could be mediated by transient low-affinity contacts poorly captured by such approaches. Consistent with this, a peptide-phage display screen suited to detect transient motif-mediated contacts identified a putative LC3-interacting motif in nsp3 (38), raising the possibility of a direct engagement. Alternatively, recruitment of LC3C, and consequently of ATG4D, to nascent DMVs may occur independently of nsp3 and nsp4 and instead be bridged by other host factors recruited to these membranes.

## Materials and Methods

### Cell culture

A549 (ATCC; ATCC-CCL-185), Calu-3 (ATCC; ATCC-HTB-55), HeLa (ATCC; ATCC-CRM-CCL-2), HEK-293T (ATCC; ATCC-CRL-3216), MRC5 (ATCC, ATCC-CCL-171), and Vero E6 (ATCC, ATCC-CRL-1586) cells were cultured in Dulbecco’s modified Eagle’s medium (DMEM, Gibco) enriched with 10% fetal bovine serum (FBS, Euroclone), 2mM L-glutamine (Sigma-Aldrich), 1mM minimal Non-Essential Amino Acids, 1mM Sodium Pyruvate (Gibco), and 10 μg/mL Penicillin/Streptomycin (Gibco). HK2 cells were cultured in DMEM/ Nutrient Mixture F-12 (Gibco) supplemented with 5% FBS, 1% insulin-transferrin-selenium, 2mM L-glutamine and 10 μg/mL Penicillin/Streptomycin. ATG4 Tetra Knockout (Cat. No. ABM-T3460) and ATG4A/B/C Stable Triple Knockout (Cat. No. ABM-T3590) HeLa cell lines were obtained from Applied Biological Materials Inc. (ABM, Richmond, BC, Canada) and were cultured in 10% fetal bovine serum (FBS, Euroclone), 25mM HEPES (Sigma-Aldrich), 2mM L-glutamine (Sigma-Aldrich), 1mM minimal Non-Essential Amino Acids, 1mM Sodium Pyruvate (Gibco), and 10 μg/mL Penicillin/Streptomycin (Gibco). All cell lines were maintained at 37 °C in a humidified incubator with 5% CO2 atmosphere. All the infections were performed in corresponding medium supplemented with 2% FBS instead of the 10% FBS. All cell lines are regularly tested for mycoplasma contamination. Knockout of ATG7 in A549-ACE2 and Calu3 cell lines was generated following the protocol of Sanjana et al.(39).

### Viruses

The SARS-CoV-2 isolate BavPat1/2020 p2, was provided by Prof. Christian Drosten through the European Virology Archive (Ref-SKU: 026 V-03883) and propagated on Vero E6-TMPRSS2 cells infected with MOI 0.001 for 72 hours at 37° C. Collected supernatant was centrifuged to remove cell debris, filtered through 0.45 um filter and supplemented with 10mM HEPES. Viral titer was determined by plaque assay on Vero E6 cells. HCoV-OC43 (ATCC, VR-1558) virus was propagated on MRC5 cells infected with MOI 0.01 for 72 hours at 37° C. Collected supernatant was cleared of cell debris by centrifugation, filtered through 0.45 um filter, and supplemented with 10mM HEPES. Viral stock was determined by TCID50 assay on MRC5 cells. Viruses were stored at −80° C. All SARS-CoV-2 infections were performed in the BSL-3 laboratory of the Telethon Institute of Genetic and Medicine (TIGEM). Decontamination of infected samples was performed according to the biosafety manual of the BSL-3 laboratory.

HCoV-OC43 and SARS-CoV-2 viral inoculum was prepared in culturing medium with reduced FBS (2%). Medium was removed from cells and viral inoculum was added on permissive cells. Cells were incubated in humidified incubator with 5% CO2 at 37°C. Viral inoculum was kept on cells for 2 hours in case of SARS-CoV-2 or 3 hours in case of HCoV-OC43 before removing it. Cells were washed 3 times with PBS and fresh 2% medium was added for the remaining time of infection.

### Lentivirus production and titration

Lentiviral stocks were produced in 70% confluent HEK 293T cells transfected with packaging plasmids pCMV-Gag-Pol and pMD2-VSV-G and backbone plasmids encoding for different genes of interest (pWPI-ACE2, pWPI-ACE2-TMPRSS2). 6 hours post-transfection the complete medium was replaced to 2% DMEM medium and 48hpt the medium containing the lentiviruses was collected cell debris removed by centrifugation and filtering through 0.45 um filter, supplemented with 10mM HEPES. Lentivirus aliquots were stored at −80°C.

Lentiviruses titration was performed in HeLa Kyoto strain, transduced with two dilutions of lentivirus preparation in presence of 4 µg/ml polybrene and then cultured for 5 days in presence of 1 µg/ml puromycin or 2 µg/ml blasticidin. The fifth day cells were washed and stained with 1% crystal violet/10% ethanol for 20 minutes. Stained cells were rinsed with water and number of colonies was used to calculate lentiviral titres.

### Generation of pBAC-SARS-CoV-2/Nluc-2A-ΔS construct

The parental pBAC-SARS-CoV-2/Nluc-2A(40) was kindly provided by Prof. Luis Martinez-Sobrido (Texas Biomedical Research Institute, San Antonio, TX, USA) to Prof. Raffaele De Francesco. To generate the S-deleted construct (pBAC-SARS-CoV-2/Nluc-2A-ΔS), the terminal region of orf1ab immediately upstream of the S open reading frame (ORF) was amplified from the parental backbone using Phusion High-Fidelity DNA Polymerase (Thermo Fisher Scientific). PCR was performed with the forward primer orf1ab-Fwd (5’-GCCTTCGAACATATCGTTTATGG-3’) and the reverse primer orf1ab-Rev (5’-CGCGGATCCTGTTCGTTTAGTTGTTAACAAG-3’), the latter introducing a BamHI restriction site. The resulting amplicon was purified using the Wizard® SV Gel and PCR Clean-Up System (Promega) and subjected to double digestion with BstBI and BamHI (New England Biolabs). This fragment was subsequently ligated into the pBAC-SARS-CoV-2/Nluc-2A backbone, which had been previously digested with the same enzymes to excise the terminal region of orf1ab and the S ORF region. The identity and integrity of the final pBAC-SARS-CoV-2/Nluc-2A-ΔS construct were confirmed by restriction enzyme analysis and Sanger sequencing.

### Cell viability analysis

Cell viability was assessed using CellTiter-Glo® Luminescent kit (Promega) according to manufacturer’s instructions. Briefly, 6,000 cells were plated in opaque-walled 96-well plates and incubated for at 37°C and 5% CO2. To measure cell viability at specific timepoints, cell culture medium was removed from each well and replaced with 100 µl of CellTiter-Glo® Reagent mixed with PBS (50% v/v). The plate was incubated at room temperature for 10 minutes to stabilize luminescent signal, which was then measured by the microplate reader (High-throughput screening microplate reader Synergy™ Neo BioTek Instruments).

### Transfection

The HeLa cells were seeded in 24-well plate with coverslips in density of 4.0 x 10^4 cells/well in complete 10% FBS medium. The following day, the medium was replaced by fresh one. Transfection mix was prepared by diluting Lipofectamine 2000 transfection reagent (Thermofisher Scientific) in reduced serum OPTIMEM medium (Thermofisher Scientific), vortexing and incubating 5 minutes at RT before adding the plasmid DNA in OPTIMEM media prepared in separate reaction tube. The ratio of 3:1 was used, corresponding to 1.5 µL transfection reagent and 0.5 µg DNA per well. After vortexing for 10 s, the mix was incubated 20 min at RT. Mixture was added to the cells drop by drop and then incubated for 30 – 48 hours.

### Nsp3-nsp4 polyprotein expression

The transfection-based nsp3-nsp4 polyprotein expression system constructs previously described by Pahmeier et al. was used(5). Transfection was performed using 1.2 µg DNA per 24-well in 1:3 ratio with Lipofectamine 2000 and incubation with cells lasted 16 hours.

### LC3C reconstitutions

Hela LC3 3xKO and control were transfected with 0,5 µg DNA per 24-well in 1:3 ratio with Lipofectamine 2000. After 4 hours, the transfection mix was removed and cells were infected with OC43 virus for another 44 hours. Reconstitution experiments with LC3C WT and mutants were performed in the OC43 system, as SARS-CoV-2 infection of HeLa cells additionally requires ectopic expression of ACE2 and TMPRSS2, which precluded achieving sufficient co-expression efficiency of the reconstituted LC3 variants under these conditions.

### Silencing using siRNA interference

Pools of three small-interfering RNAs targeting different autophagy factors together with the positive control ACE2 and a non-targeting control were acquired from Dharmacon. For further validation of ATG4D KD effect, two individual siRNA targeting ATG4D, ACE2 and non-targeting control siRNA were purchased from Ambion (Thermoscientific) and pooled together. 5x concentrated mastermix (100 nM) for silencing was prepared by adding siRNA pool to OPTIMEM (mixture A) and OPTIMEM containing the Lipofectamine RNAiMAX (ThermoFisher) transfection reagent (mixture B). Both mixtures were added together, homogenised by pipetting and incubated 30 min at RT. Then the 5x concentrated master mix was added to cells in 10% complete medium to achieve 1x concentration (20nM siRNA). Cells were incubated in humidified incubator with 5% CO2 for 48 hours.

### Immunofluorescence

Infected and mock A549 cells were grown on glass coverslips with density 50.000 cells/well in 24-well plate format. At specific times post-infection cells were fixed with 4% EM grade paraformaldehyde (PFA) in PBS at room temperature for 30 min. Then, whole plates were submerged in 4% PFA for another 30 minutes to be taken out of BSL-3 laboratory. PFA was removed and cells washed with PBS 2x. The permeabilization and blocking of cells was performed in saponin buffer (0.05% saponin, 0.5% BSA, and 50 mM NH4Cl in PBS) for 30 minutes. Coverslips were incubated with the primary antibodies at 4°C overnight or room temperature (RT) for 2 hours. Coverslips were washed with PBS-0.01% Tween-20 and incubated with secondary antibodies for 1 hour at RT. Cells were then washed with PBS-0.01% Tween-20, stained for 15 minutes with Hoechst 33342, washed with ddH2O and mounted on the microscope slides using ProLong™ Glass Antifade Mountant™. The antibodies used in this study are reported in Table 1.

**Table 1.** List of antibodies used for immunofluorescence.

| Antibody anti- | Origin | Dilution | Manufacturer | Identifier Cat.# |
| --- | --- | --- | --- | --- |
| SARS-CoV-2 N | Mouse | 1:1000 | Sino Biological | 40143-MM05 |
| SARS-CoV-2 nsp3 | Rabbit | 1:1000 | Genetex | GTX135589 |
| HCoV-OC43 N | Rabbit | 1:2000 | Sino Biological | 40643-T62 |
| dsRNA | Mouse | 1:500 | Scicons | 10010500 |
| LC3B | Rabbit | 1:500 | MBL | PM036 |
| HA | Rabbit | 1:400 | Sigma-Aldrich | H6908100UL |
| V5 | Mouse | 1:500 | Sino Biological | 100378-MM04 |
| Alexa Fluor Plus 488 anti-mouse IgG (H+L) | Goat | 1:800 | Invitrogen | A32723 |
| Alexa Fluor 488 anti-mouse IgG1 | Goat | 1:800 | Invitrogen | A21121 |
| Alexa Fluor 488 anti-mouse IgG2A | Goat | 1:800 | Invitrogen | A21131 |
| Alexa Fluor 488 anti-mouse IgG2B | Goat | 1:800 | Invitrogen | A21134 |
| Alexa Fluor 488 anti-rabbit | Goat | 1:800 | Invitrogen | A32731 |
| Alexa Fluor Plus 568 anti-mouse IgG (H+L) | Goat | 1:800 | Invitrogen | A11031 |
| Alexa Fluor 568 anti-mouse IgG1 | Goat | 1:800 | Invitrogen | A21124 |
| Alexa Fluor 568 anti-mouse IgG2A | Goat | 1:800 | Invitrogen | A21141 |
| Alexa Fluor 568 anti-mouse IgG2B | Goat | 1:800 | Invitrogen | A21144 |
| Alexa Fluor 568 anti-rabbit | Goat | 1:800 | Invitrogen | A11011 |
| Alexa Fluor 647 anti-mouse | Goat | 1:800 | Invitrogen | A11011 |
| Alexa Fluor 647 anti-rabbit | Goat | 1:800 | Invitrogen | A32733 |

#### qRT-PCR

The total RNA was extracted using Maxwell® RSC simply RNA Cells Kit (Promega) according to the manufacturer’s protocol. cDNA was synthesized from 200-400 ng of RNA with LunaScript RT SuperMix Kit according to manufacturer’s instruction. A 1:20 dilution of the cDNA was used directly for qPCR analysis using specific primers (primer sequences in Table 2) and PowerUp SYBR Green Master Mix for qPCR (Applied Biosystems). The mRNA relative abundance from each sample was evaluated based on its cycle threshold (ct) values normalized to hypoxanthine phosphoribosyltransferase 1 (HPRT) transcript levels.

**Table 2.**
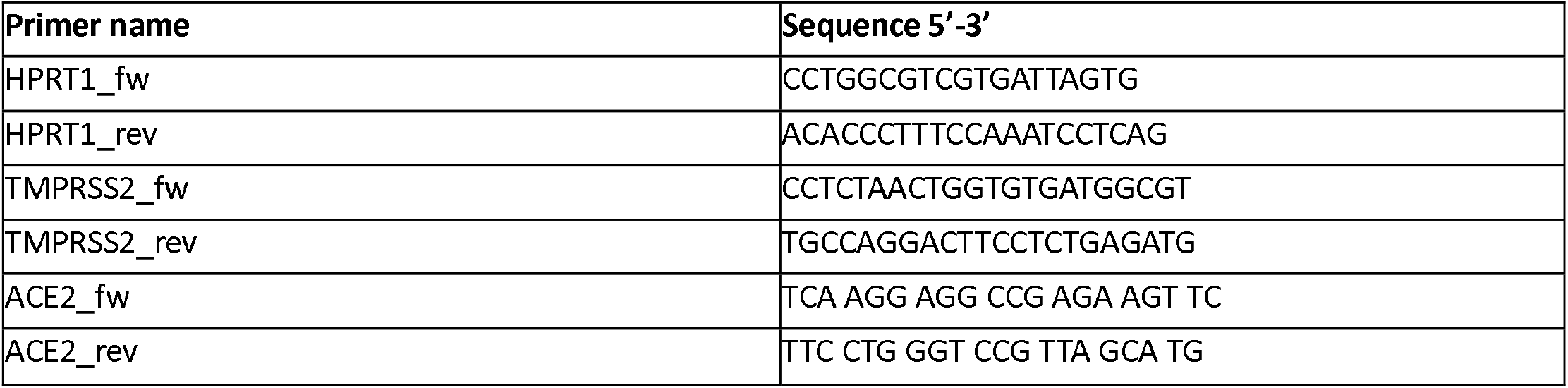
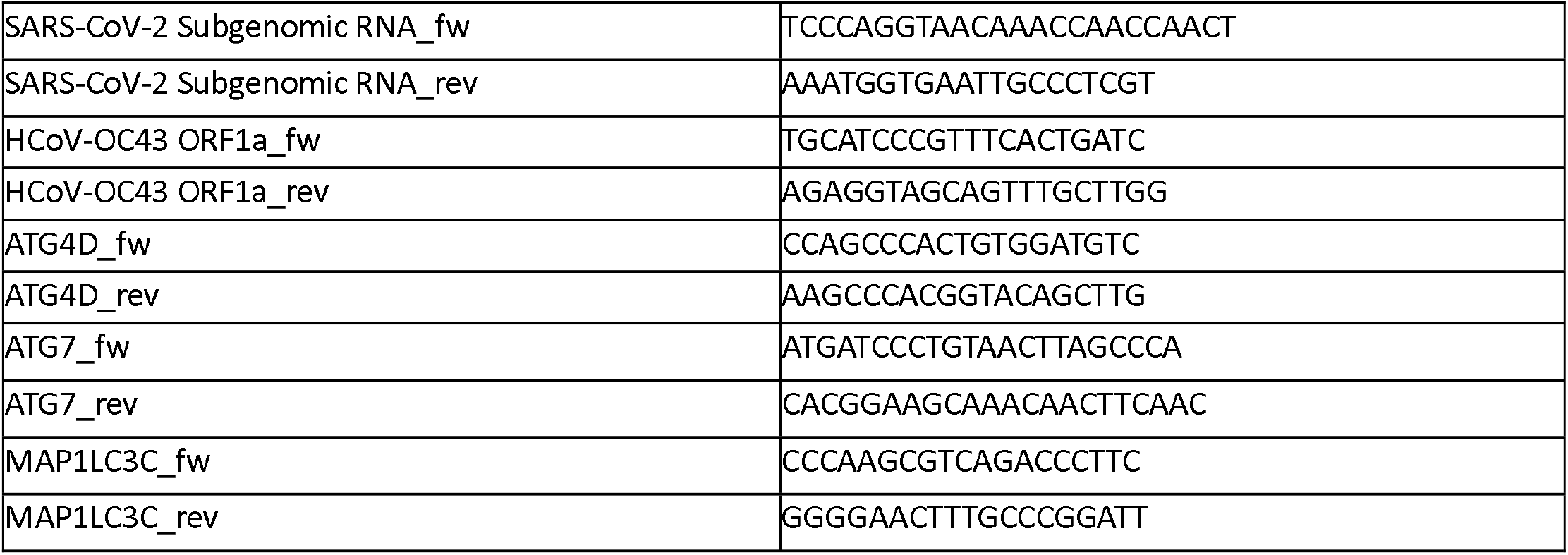
list of primers used for qPCR.

#### Western Blot

Cells were washed with PBS and lysed in RIPA lysis buffer supplemented with protease inhibitor cocktail (Complete EDTA-free, Sigma-Aldrich), incubated 20 minutes on ice and centrifuged at 20000 g for 10 minutes in table-top centrifuge. Protein amount in the sample was quantified Bradford Protein Measurement Assay (BioRad). 20-25 ug of protein was mixed with 6x sample buffer (375 mM Tris-HCl at pH 6.8, 12% SDS, 60% glycerol, 0.6 M DTT, 0.06% bromophenol blue) to achieve 1x concentration and denaturated for 5 minutes at 95°C. Total protein was resolved by homemade SDS-PAGE, mPAGE™ 4-12% or 4-20% Bis-Tris Precast Gel (Sigma-Aldrich) MOPS running buffer (Thermofisher) and transferred to polyvinylidene fluoride (PVDF) 0.45 µm membrane (0.2 µm PVDF membrane for LC3B blotting) for 90 minutes at constant current – 250 mA for all proteins except of LC3B, which was transferred at 200 mA. Membranes were blocked with 3% BSA or 5% skimmed milk in TBS-T (TBS with 0.1% Tween-20) for 1 hour at room temperature. Incubation with primary antibodies was performed overnight at 4°C. Used antibodies are listed in Table 3. Membranes were washed three times with TBS-T and incubated for 1 hour with horseradish peroxidase (HRP)-conjugated secondary antibodies in TBS-Tween (0.1%). Chemiluminescent signal was captured using the ECL system. Image processing and band analysis was performed using Fiji software (NIH).

**Table 3.** List of antibodies used for western blot.

| Antibody anti- | Origin | Dilution | Manufacturer | Identifier Cat.# |
| --- | --- | --- | --- | --- |
| GAPDH | Mouse | 1:5000 | Santa Cruz Biotechnology | sc365062 |
| vinculin | Mouse | 1:5000 | Santa Cruz Biotechnology | sc-73614 |
| β-actin | Mouse | 1:5000 | Sigma-Aldrich | A5441 |
| LC3B | Mouse | 1:1000 | Nanotools | NTL0231100 |
| HA | Mouse | 1:1000 | Biolegend | 901501 |
| ATG7 | Rabbit | 1:1000 | Cell Signaling | 8558 |
| p62 (SQSTM1) | Mouse | 1:1000 | Cell Signaling | 88588 |
| N (SARS-CoV-2) | Rabbit | 1:1000 | Cell Signaling | 26369 |
| M (SARS-CoV-2) | Rabbit | 1:500 | Kindly provided by Carolyn Machamer |  |
| ACE2 | Mouse | 1:250 | R&D Biotechne | MAB9332-100 |
| anti Rabbit IgG (whole molecule) peroxidase | Goat | 1:6000 | Sigma-Aldrich | A6154 |
| anti Mouse IgG (whole molecule) peroxidase | Goat | 1:6000 | Sigma-Aldrich | A4416 |

#### Pellet embedding

Cells grown in 6-well plate format were washed once with PBS and then fixed with 1% glutaraldehyde (GA) in 0.2 M HEPES (pH-7.3!) buffer 0.5 ml/12well for 30min at RT. Then washed once with PBS and wells were filled with 4% EM grade PFA for another 30 minutes to exit the BSL-3 laboratory. Afterwards, 3 washes with PBS were performed and cells were overlayed with 500 ml/well of 1% BSA diluted in PBS 1X. Cells were scraped and centrifuged at 13200rpm for 10 minutes, to obtain cell pellet. Post-fixation was performed by incubation with a mixture of 2% OsO4 and 3% potassium ferrocyanide on ice for 30 minutes. Pellet was washed three times with ddH_2_O, incubated for 5 minutes with 1% thiocarbohydrizide diluted in H_2_O, washed again and treated with 0.5% uranyl acetate overnight at 4°C. After the incubation pellets were rinsed with water, progressively dehydrated with increasing concentrations of ethanol (50% to 100%) and treated with 100% acetone. Then the mixture of acetone and epon was added and cells subsequently embedded in a polymerizing Epon for 48 h at 60°C. Embedded cells were cut in 70-nm thick sections by using a Leica UC7 microtome and a diamond knife (Diatome).

#### Electron microscopy

Thin and thick sections (70 nm and 800 nm, respectively) were decompressed with chloroform and collected onto formvar carbon slot grids (Prod No. 01805-F, Ted Pella). Thin sections were imaged with a Tecnai Spirit 120 kV. Electron tomography was performed on thick sections at the National Facility for Structural Biology – IU1 Cryo-Electron Microscopy, Fondazione Human Technopole, Milan, Italy. Tilt series were acquired on a Thermo Scientific Spectra 300 in STEM mode with the HAADF detector, in SerialEM, tilt range from +/-65 with 1.5 degrees increment. The tilt series were reconstructed on IMOD (version 5.1.9). Segmentation and 3D reconstruction for this paper was generated using Dragonfly software, Version 2022.2 for Windows (Comet Technologies Canada Inc., Available from: https://dragonfly.comet.tech/. Accessed 2025 Oct 2).

#### Bioinformatic analysis

Images were analysed with the Fiji image analysis software. Graphs and statistical analysis were performed using GraphPad Prism software. Schemes were created in BioRender. Image panels were designed in Adobe Illustrator. Statistical significance was assessed using a one-sample t-test against comparing experimental values with the theoretical mean of 1 of control cells (ns [not significant] p>⍰0.05, *p<⍰0.05, **p⍰<⍰0.01, ***p⍰<⍰0.001, **** p⍰<⍰0.0001).

## Data availability

All data generated or analyzed during this study are included in this article and its supplementary information files. Requests for resources and reagents should be directed to and will be fulfilled by the lead contact, Mirko Cortese.

## Acknowledgments

We thank Sofia Maria Luigia Tiano, Federica Camerota, Debora Stelitano and Gennaro Bianco for technical support, constructive discussions and collaborative environment throughout the course of this work.

All the schematic figures were generated with Biorender. We thank the High Content Screening facility, the Advanced Microscopy and Imaging facility and the Mass Spectrometry facility of the TIGEM for their support. We acknowledge the Access and services provided by the National Facility for Structural Biology – IU1 Cryo-Electron Microscopy, Fondazione Human Technopole, Milan, Italy; Call for Access 25-ROUND-1, Project ID2166684.

MC is supported by the Human Technopole Early Career Fellowship Programme (HT-ECF 763 Programme). ŠV was supported by a PhD Fellowship from the Scuola Superiore Meridionale (SSM), University of Federico II. Work of the Herhaus lab at the Helmholtz-Zentrum für Infektionsforschung GmbH, Braunschweig is supported by the Microbial Stargazing program of the German Federal Ministry of Education and Research (BMFTR) under grant number 01KX2324 and the CRC project on selective autophagy: project-ID 259130777. Responsibility for the content of this publication lies with the author.

## Author contributions

ŠV and LI designed/performed most of the experiments and analysed the data, VM performed some experiments and data analysis, EP performed TEM samples acquisitions, LD, RdF, PG, CS, MCi, LH provided material and experimental support, MC and ŠV conceived the study; MC and LH managed the project and supervised experiments, ŠV, MC and LH wrote the manuscript with input from all authors.

## Materials & Correspondence

For any requests concerning material generated in this study, contact the corresponding authors

## Figure Legends

**Supplementary Figure 1: LC3B accumulates upon SARS-CoV-2 infection**. (**a**) SDS-PAGE and immunoblot analysis of A549-ACE2-TMPRSS2 cells using antibodies to the SQSTM1, SARS-CoV-2 N and M proteins and LC3B. GAPDH was used as a loading control. Cells were collected at 24-, 48- and 72-hours post-infection prior to lysis. Uninfected cells served as control. (**b**) Representative image of LC3B puncta accumulation in infected 549-ACE2-TMPRSS2 cells. The cells were infected with SARS-CoV-2 for 24 hours, permeabilised and stained for N protein (yellow) and LC3B (grey). Scale bars: 20⍰μm. Yellow arrows indicate accumulation of LC3 puncta in infected cells. Dashed line indicates mock cells. (**c**) Quantification of LC3B puncta in mock and infected cells. Each dot represents one cell (N=100). (**d**) HeLa 3xKO LC3 and control cells were seeded in 96 well plates and 24, 48 and 72 hours post seeding, the cells were lysed and total ATP levels were measured to assess cell growth and viability by luminescence. (**e**) HeLa 3xLC3 KO and their control cells were transfected with nsp3-GFP plasmid to evaluate translation capacity based on GFP intensity. Statistical significance was determined using t-test (ns [not significant], ****p⍰<⍰0.0001).

**Supplementary Figure 2: Characterization of ATG7 KO** (**a**) Representative images of A549-ACE2-ATG7 KO and WT cells upon EBSS-mediated autophagy induction. Cells were infected incubated with EBSS medium for 4 hours and then fixed. The cells were permeabilised and stained for LC3B (white). Scale bars: 20⍰μm. (**b**) Quantification of LC3B puncta number in ATG7 KO and control cells. Data shown in graphs are mean ± s.e.m. from 3 independent experiments, each performed in technical replicates. Statistical significance was assessed using t-test. (**** p⍰<⍰0.0001).

**Supplementary Figure 3: Characterization of ATG4D in HK2 and OE cells** (**a**) HK2 cells were incubated for 48h hours with 20nM siRNA targeting ATG4D, ACE2 (positive control) or non-target (NT, negative control). Cells were then infected with SARS-CoV-2 moi=1 for 24 hours. RNA was harvested, and q-RT-PCR was used to evaluate the viral replication based on subgenomic mRNA relative levels. (**b**) The number of SARS-CoV-2-positive HK2 cells was quantified in 10 fields of view for each condition. (**c**) Extracellular infectivity of siRNA targeted HK2 cells was determined using a plaque assay in Vero E6 cells (**d**) HK2 cells were transfected with 20nM siRNA and lysed 48 hours post transfection. The ACE2 and LC3B proteins were detected by western blot. GAPDH served as loading control. (**e**) HeLa WT, ATG4 triple KO (A/B/C) and ATG4 tetra KO (A/B/C/D) were transfected with nsp3-GFP plasmid to evaluate translation capacity based on GFP intensity. (**f**) Schematic representation of pWPI lentiviral constructs containing the HA-tagged ATG4D isoforms. (**g**) SDS-PAGE and immunoblot analysis of A549 cells transduced with different moi (1, 10, 20) of lentiviruses (either empty construct, ATG4D isoform1 or ATG4D isoform 2). Cells were lysed in 6-well format and HA tag protein was detected by western blot. Vinculin served as loading control; protein molecular mass in kilodalton is given on the right. Quantification of the ratio between HA and vinculin is shown in the right panel. Black arrowheads show conditions which were selected for further experiments. (**h**) Representative image of ATG4D localisation in A549 cells with HA tagged ATG4D isoform 1 OE. The cells were permeabilised and stained for HA protein (white), TOMM20 (red). Nuclei were visualised by Hoechst staining. (**i**) Representative image of ATG4D localisation in A549 cells with HA tagged ATG4D isoform 2 OE. The cells were permeabilised and stained for HA protein (white), TOMM20 (red). Nuclei were visualised by Hoechst staining. (**j**) Representative images of SARS-CoV-2 infected A549 cells overexpressing HA tagged ATG4D isoform 2. Cells were infected with moi=1 of SARS-CoV-2 and fixed 24hpi. The cells were permeabilised and stained for HA (red) and dsRNA(green) and TOMM20 (blue). Scale bars: 5⍰μm. Data shown in a-c are mean ± s.e.m. from 3 independent experiments, each performed in technical triplicates. Statistical significance was assessed using a one-sample t-test against comparing experimental values to NT with the theoretical mean of 1 (ns [not significant] p⍰>⍰0.05, *p⍰<⍰0.05, **p⍰<⍰0.01, ***p⍰<⍰0.001, **** p⍰<⍰0.0001).

**Supplementary Figure 4: LC3C expression control** (**a**) U2OS-eGFP-mCherry-LC3C WT and LC3C S93/96A cells (top and bottom panels, respectively) were sorted for eGFP and mCherry signal in FACSAriaII SORP cell sorter to achieve comparable LC3C expression levels. Gating strategy for the two cell lines is reported.

