## Supplementary figures and images for "LC3C–ATG4D regulatory axis supports coronavirus replication organelle formation independently of canonical autophagy"

### Supplemental Figures

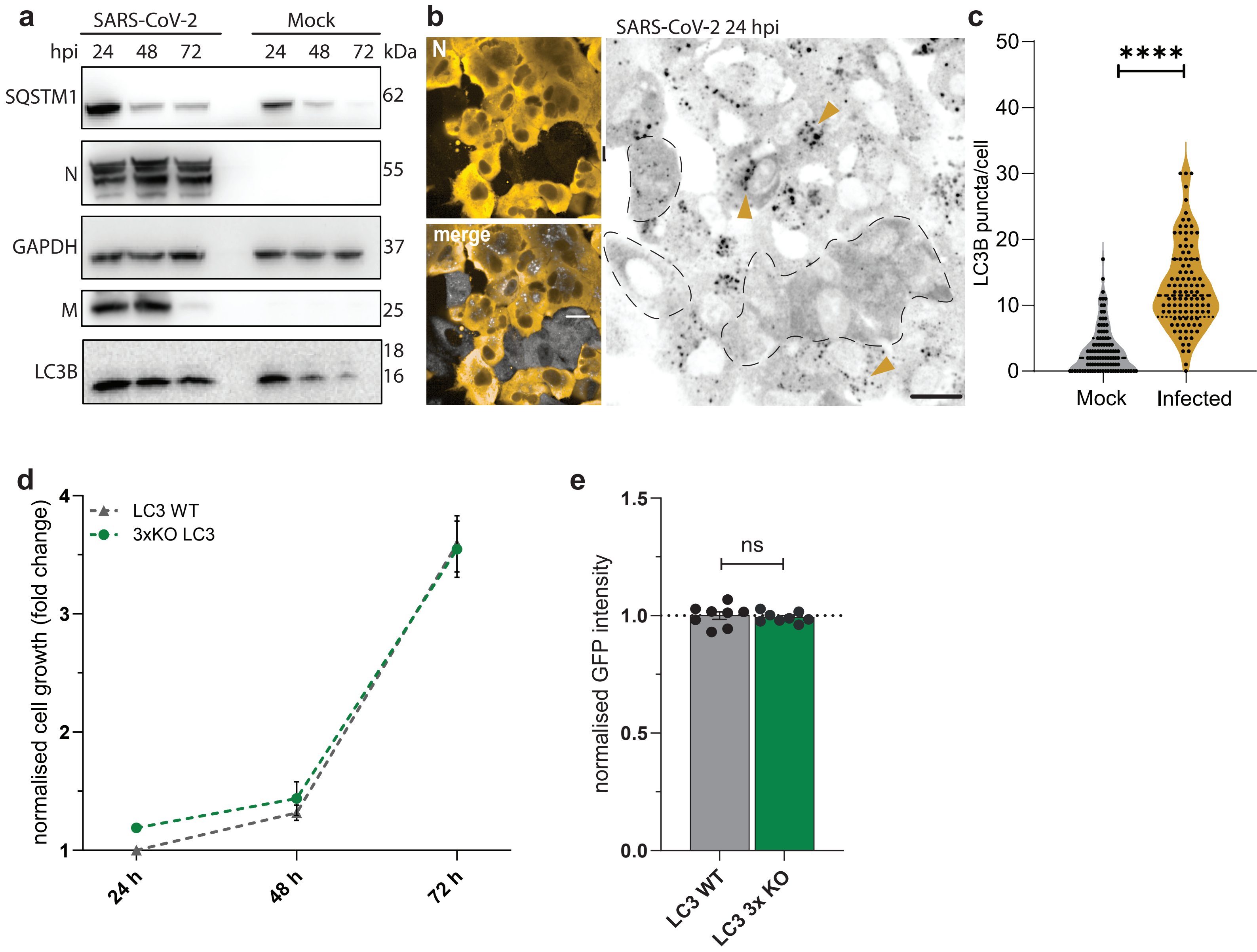

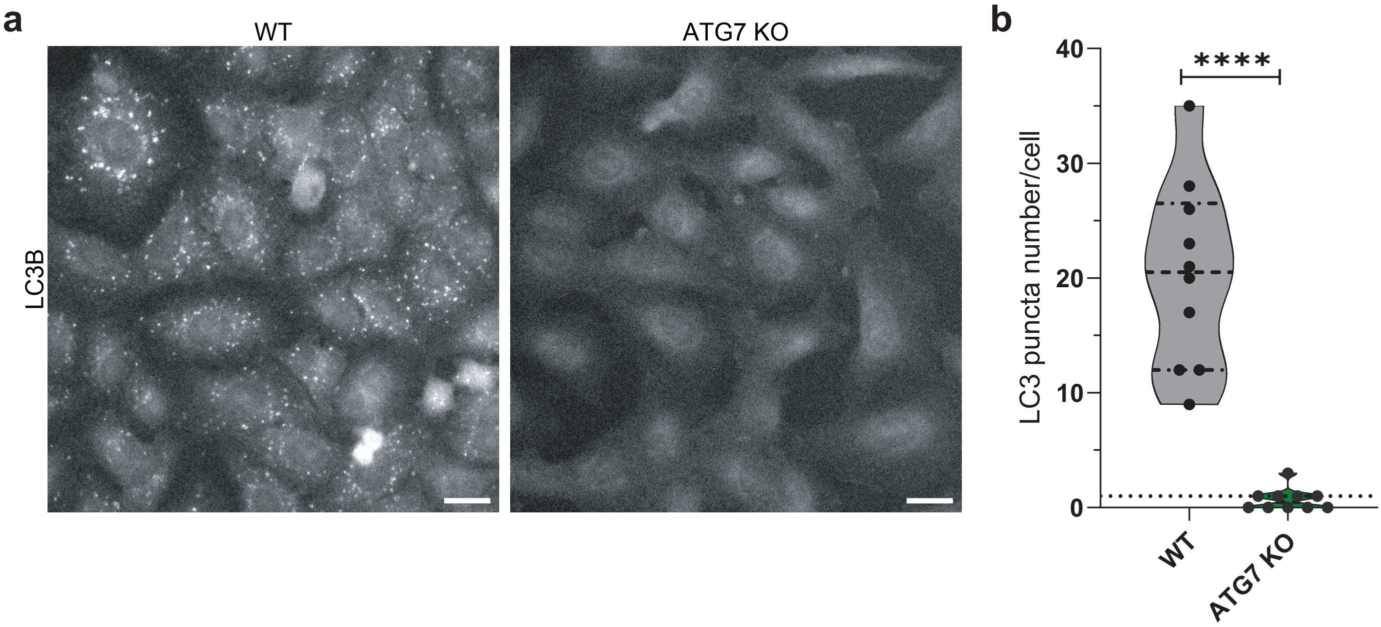

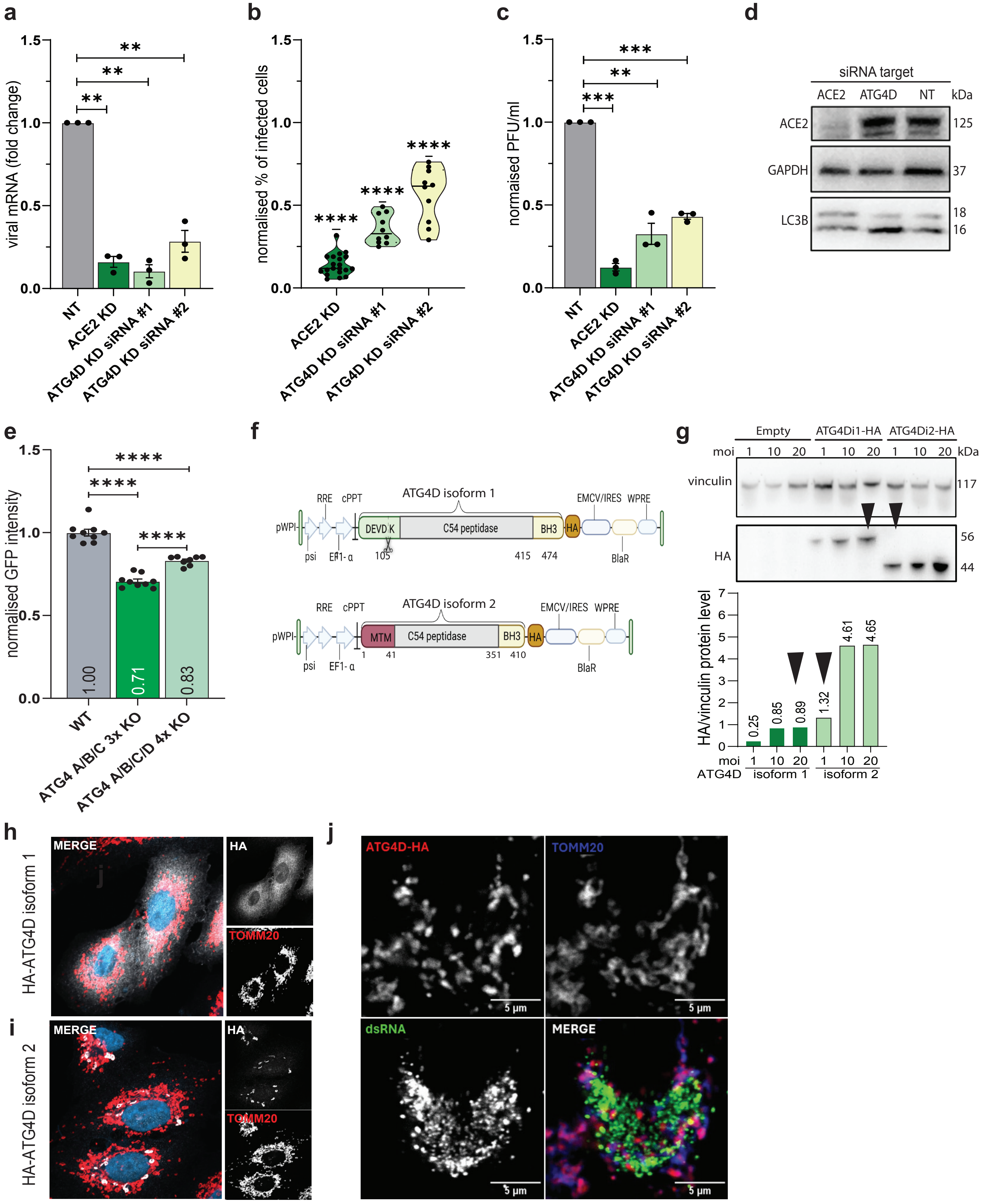

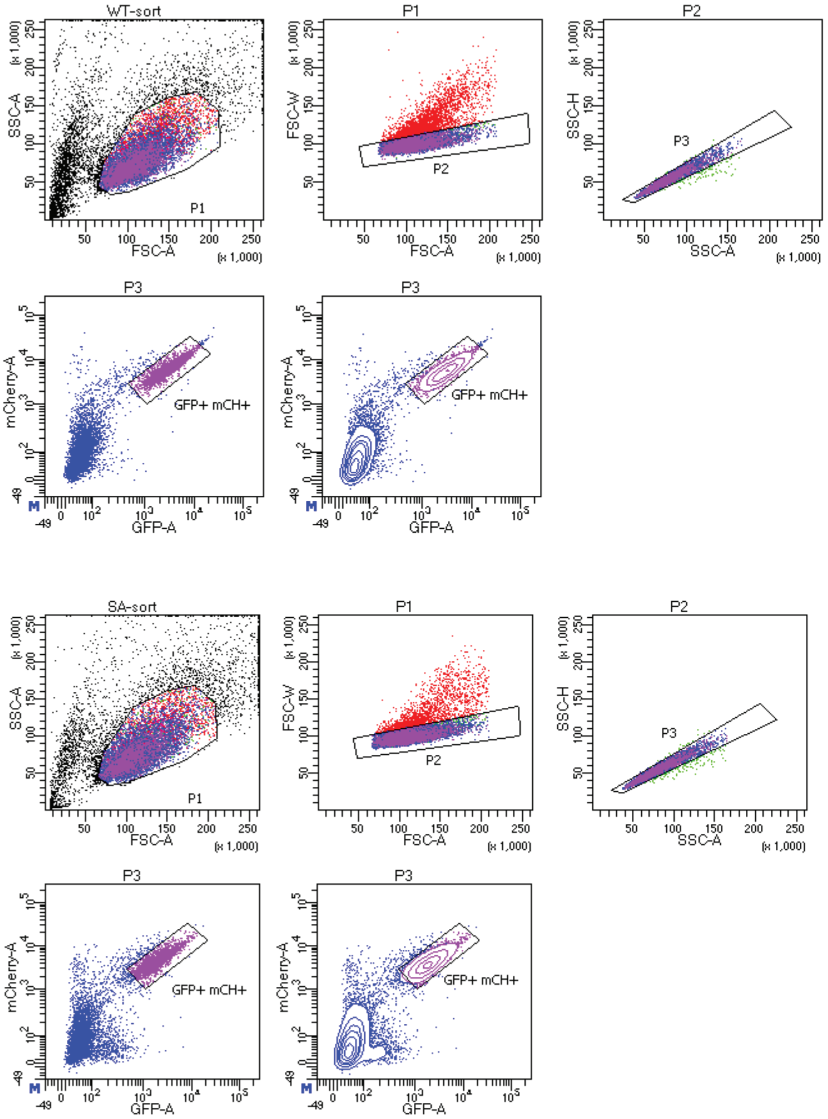
